# Tissue-specific responses of the central carbon metabolism in tomato fruit to low oxygen stress

**DOI:** 10.1101/2024.10.31.621370

**Authors:** Xindan Li, Konstantinos Terzoudis, Maarten L.A.T.M. Hertog, Bart M. Nicolaï

## Abstract

Tomato (*Solanum lycopersicum* L.) is an important model plant whose fleshy fruit consists of well-differentiated tissues. Recently it was shown that these tissues develop hypoxia during fruit development and ripening. Therefore, we employed a combination of metabolomics and isotopic labeling to investigate the central carbon metabolic response of tomato fruit tissues (columella, septa and mesocarp) to low O_2_ stress. The concentration and ^13^C-label enrichment of intermediates from the central carbon metabolism were analyzed using gas chromatography–mass spectrometry. The results showed an increase in glycolytic activity and the initiation of fermentation in response to low O_2_ conditions. In addition, the up-regulation of the GABA shunt and accumulation of amino acids, alanine and glycine, were observed under low O_2_ conditions. Notably, tissue specificity was observed at the metabolite level, with concentrations of most metabolites being highest in columella tissue. In addition, there were tissue-specific differences in the central carbon metabolism with the columella exhibiting the highest metabolic activity and sensitivity to the changes in O_2_ concentration, followed by septa and mesocarp tissues. Our results are consistent with common plant responses and adaptive mechanisms to low O_2_ stress, while unravelling some tissue-specific differences, increasing our understanding of the intact fruit response to low O_2_ stress.

## 1. Introduction

Tomato (*Solanum lycopersicum* L.) has been widely used as a model system to study fleshy fruit (Pabón-Mora and Litt, 2011). The tomato fruit is composed of several distinct tissues: the outer pericarp, which includes the exocarp, mesocarp, and endocarp, as well as the radial pericarp (septa), which separates the ovary in which the seeds are located. These seeds are protected by the surrounding locular gel and are connected to the internal placenta. In addition, the fruit has a central columella structure (Chamley *et al*., 2019). 3D microstructural analysis of tomato fruit tissues using X-ray computed tomography showed that mesocarp and septa tissue possess larger cells but smaller and less interconnected pores as compared to the placenta and columella (Xiao *et al*., 2021). The complex anatomy of tomato fruit with its distinct tissues refers to their functional diversification and specialization (Shinozaki *et al*., 2018). The differentiation of tissue has been demonstrated in some targeted and large-scale omics studies, concerning ethylene biosynthesis (Van de Poel *et al*., 2014), photosynthesis (Smillie *et al*., 1999), volatile biosynthesis (Wang *et al*., 2018), sugar metabolism (Schaffer and Petreikov, 1997), their transcriptional profile (Shinozaki et al., 2018; Matas et al., 2011) and their metabolomic profile (Moco *et al*., 2007; Chamley *et al*., 2019). However, most tomato studies have focused on the outer pericarp tissue, with few research addressing the physiology and biochemistry of the inner tissues (Van de Poel *et al*., 2014).

The central carbon metabolism is an important biochemical process that enables cells to survive and grow by providing energy and intermediates for other metabolic pathways. The central carbon metabolism includes key biochemical pathways such as the glycolysis, tricarboxylic acid (TCA) cycle, mitochondrial electron transport chain (mETC), pentose phosphate pathway (PPP), and fermentation (Boeckx *et al*., 2019). The central carbon metabolism in tomato fruit has been extensively researched using intact fruit or pericarp tissue to understand its key role in fruit development, ripening, and postharvest quality (Oms-Oliu *et al*., 2011; Colombié *et al*., 2017; Quinet *et al*., 2019). O_2_ is the final electron acceptor in the mETC and directly influences energy production in the central carbon metabolism (Ho et al., 2013). When there is limited or no O_2_ available inside the fruit, the tissues are subjected to low O_2_ stress, which triggers a major shift in metabolism from aerobic respiration to fermentation, leading to a depletion of carbon reserves and causing off-flavors in the fruit (Geigenberger, 2003; António *et al*., 2016; Boeckx *et al*., 2019). Xiao et al. (2024) reported for tomato fruit at different ripening stages that under atmospheric conditions (20 °C), the O_2_ concentration in the locular gel goes down to around 1-7 kPa, while that of the other tissues typically is about 8-17 kPa. During growth, tomatoes are typically exposed to field or greenhouse temperatures well above 20 °C. Ho et al. (2018) have shown that elevated temperatures result in increasingly lower O_2_ concentrations within fruit tissues. This implies that the fruit internal O_2_ levels may be even lower than those reported by Xiao et al. (2024) at 20 °C. In addition, Ye et al. (2015) found that some fermentation-related genes were significantly up-regulated in pericarp tissue of tomato fruit cultivated under a diurnal temperature regime of 28/20 °C, indicating low O_2_ stress inside the fruit’s pericarp. However, tissue-specific responses and regulatory mechanisms of the central carbon metabolism to low O_2_ stress in tomato fruit are not understood well.

Metabolomics is instrumental in providing a comprehensive view of metabolites and their interactions, thus illuminating biochemical pathways and aiding in understanding organism-environment relationships at the molecular level (Brunetti *et al*., 2013; Aizat *et al*., 2018). However, understanding metabolite levels alone does not unveil the actual fluxes through pathways, nor the possible regulatory mechanisms (Fernie *et al*., 2005). To fully grasp the control and regulation of metabolic networks, it is necessary to introduce isotope-labeled substrates into the metabolism (Ratcliffe and Shachar-Hill, 2006; Ampofo-Asiama *et al*., 2014). Feeding whole tomato fruit with isotope-labeled substrate is challenging, as it is difficult to ensure that the label is evenly distributed to all tissues. To overcome this problem, we used tissue discs, allowing for more precise control over substrate exposure and ensuring consistent labeling across samples. Since tomato are climacteric fruit, they continue to ripen during the experimental process as long as the fruit has developed fully. Additionally, the process of collecting tissue discs causes wounding, which will accelerates ripening (Boller and Kende, 1980). Consequently, this makes it difficult to study intermediate ripening stages as they cannot be assumed stable enough over time. Therefore, we focused on the red ripened stage, where the fruit is fully matured and the metabolic processes are more stable for analysis.

The aim of this work was to investigate possible tissue-specific responses and adaptive mechanisms of the central carbon metabolism to low O_2_ stress in red ripe tomato fruit. Our work focused on columella, septa, and mesocarp tissue given their different positions within the tomato fruit structure. The locular gel and placenta were omitted due to the practical challenges of collecting such tissue from red ripe tomato fruit. The central carbon metabolism primarily involves the formation and breakdown of carbon-carbon bonds, with glucose serving as the initial substrate to glycolysis. Therefore, ^13^C_6_ glucose was utilized as a labeled substrate to feed the tissues. Fresh tissues (columella, septa and mesocarp) were excised from intact fruit and incubated in a uniformly labeled ^13^C_6_ glucose buffer aerated with either 21, 1, or 0 kPa of O_2_. The concentration and ^13^C-label enrichment of intermediates in the central carbon metabolism were analyzed using GC–MS.

## 2. Materials and methods

### 2.1. Fruit material

Tomato fruit (*Solanum lycopersicum* L. ‘Mattinaro’) were harvested at the red ripe stage for experiments. For the feeding experiment, tomato fruit were sourced in August 2022 from a commercial greenhouse in Herenthout, Belgium (51°07’57.7”N 4°43’55.8”E). Tomato fruit were disinfected with 2 % sodium hypochlorite solution for 2 min and then washed with tap water before being used in the feeding experiment. Tomato fruit for tissue respiration experiment were obtained in June 2022 from the experimental station for vegetable cultivation in Sint-Katelijne-Waver, Belgium (51°04’39.4”N 4°31’38.5”E).

### 2.2. Osmolality of different tissues measurement

The osmolality of individual columella, septa and mesocarp tissues was measured using the method by Beshir et al. (2017) using an Osmomat 3000 basic device (Gonotec, Berlin, Germany). The molarity of the medium used during the feeding experiments of the three tissues was adjusted accordingly (See Supplementary Table S1 for the exact values).

### 2.3. Feeding experiment

The feeding experiment was conducted using ^13^C_6_ glucose as the labeled carbon substrate, following the procedure described by Beshir et al. (2017). Equatorial slices (3-4 mm thick) were cut from the fruit using a meat slicer. A cork borer (∅ 9 mm) was used to separate the slice into the different tissues (columella, septa, and mesocarp) by cutting out smaller tissue discs from these fruit slices. This approach guaranteed a constant surface area for labeled substrate uptake, facilitating the incorporation of quantifiable label amounts into the metabolite pools within a short timeframe. Per tissue, the collected discs were washed to remove damaged cells using an isotonic solution containing 2 mmol L^−1^ CaCl_2_, 2 mmol L^−1^ MgCl_2_, 2 mmol L^−1^ DTT (1,4-dithiothreitol), 50 mmol L^−1^ HEPES/KOH buffer (pH 7.0) with betaine used as an osmoticum. After washing, 6 g of tissue discs were placed in separate flasks each containing 15 mL buffer solution identical to the one used for washing the tissue discs, but supplemented with 20 mmol L^−1^ unlabeled glucose, and continuously aerated with 21 kPa O_2_ at 18 °C (15 L h^−1^). A preincubation period was set to overcome the initial impact of handling and wounding (Boeckx, 2018).

After pre-incubation in an unlabeled solution for 24 h (Supplementary Method S1), the tissue discs were transferred to a solution of the same composition containing 20 mmol L^−1^ ^13^C_6_ glucose (≥99 % enrichment). Throughout the experiment, the buffer solution was continuously aerated with humidified air with 15 L h^−1^ at 18 °C containing 21, 1 or 0 kPa O_2_ balanced by N_2_. Tissue and buffer samples were collected in triplicate at different time points (0 h, 2 h, 4 h, 8 h, 12 h and 24 h). At the same time points, gas samples were obtained from the outlet of the experimental set-up using 1 L fluorinated ethylene propylene (FEP) bags with polypropylene valves (Sense 147 Trading, Groningen, The Netherlands). After collection, the tissue samples were washed three times with a hypotonic solution of 50 mmol L^−1^ HEPES/KOH buffer (pH 7.0) to remove excess labeled substrate and medium salts. Subsequently, the samples were rapidly frozen in liquid nitrogen, ground into powder, and stored at −80 °C for metabolite analysis.

### 2.4. Respiration measurements of different tissues

The respiration rates of the three tissues were measured according to the method described by Ho et al. (2008) with some modifications. Columella, septa and mesocarp tissue discs were prepared in the same way as the samples from the feeding experiment. The tissue discs were placed into the petri dish and then into a 1.7 L glass jar along with polyvinyl chloride (PVC) cylindrical block (volume of about 0.5 L) to reduce the free space in each jar. Five different gas conditions were used in this experiment including 21, 10, 3, 1 and 0 kPa of O_2_ with 0 kPa of CO_2_. For each gas condition, the jars were flushed at 18 °C for 20 min with the specified gas mixture. After flushing, the valves of each jar were closed, and the headspace O_2_ and CO_2_ concentrations of each jar were measured initially and after 16 h using a gas analyzer (Checkmate III, PBI Dansensor, Denmark). The headspace total pressure was recorded using a pressure sensor DPI 142 (GE Druck, Germany).

The Michaelis-Menten equation was used to determine the reaction rates of O_2_ and CO_2_ in the three tissues based on the experimental data of O_2_ consumption rate and CO_2_ production rate (nmol kg^−1^ s^−1^ FW) (Hertog *et al*., 1998):

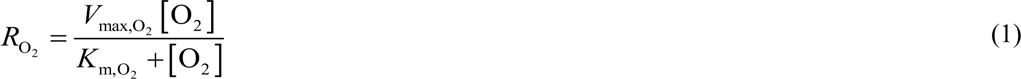

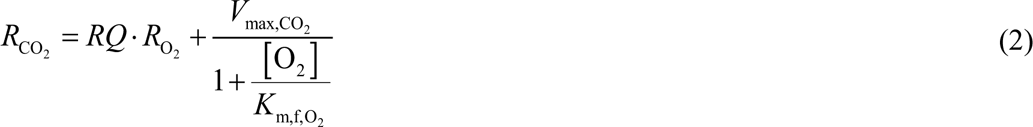

where [O_2_] is the O_2_ concentration (kPa); *V*_max,O2_ (nmol kg^−1^ s^−1^) is the maximal O_2_ consumption rate; *RQ* is the respiratory quotient; *V*_max,CO2,_ (nmol kg^−1^ s^−1^) is the maximal fermentative CO_2_ production rate; *K*_m,O2_ is the Michaelis constant for O_2_ consumption (kPa); *K*_m,f,O2_ is the Michaelis constant for the competitive inhibition of fermentative CO_2_ production by O_2_ (kPa).

### 2.5. Analysis of fermentation metabolites (ethanol and acetic acid)

The concentration of ethanol and acetic acid was measured using the commercial enzymatic kits K-ETOH and K-ACETRM (Megazyme, Wicklow, Ireland), respectively. For tissue samples, a 200 mg sample was placed in an Eppendorf tube along with 500 µL of distilled water. The mixture was then incubated at 60 °C for 30 min. After cooling it to 4 °C for 10 min, the sample was centrifuged at 22,000 g for 20 min at 4 °C, and then left at 4 °C for 1 h before analysis. Finally, 200 µL of supernatant was used for further analysis following the instructions of the microplate assay procedure. For buffer samples, 1 mL of buffer was centrifuged at 22,000 g for 20 min at 4 °C. Of the supernatant, 200 µL was analyzed as described in the microplate assay procedure. Measurements of volatile ethanol and ethyl acetate as collected into the sampling bags were carried out using a Voice200ultra® SIFT-MS instrument (Syft Technologies, Christchurch, New Zealand). The Selected-Ion Flow-Tube Mass Spectrometer (SIFT-MS) used helium as a carrier gas with a flow of 320.69 mL s^−1^ and the sample was introduced into the system with a flow of 21 mL min^−1^. A Selected Ion Mode (SIM) scan was used to measure the samples with a scan time of 120 ms and with a time limit of 100 ms and a count limit of 1000 counts, utilizing the reagent ions H_3_O^+^, NO^+^ and O_2_^+^. When analyzing ethyl acetate, specific product ions were selected, including H_3_O^+^ at a mass-to-charge (m/z) ratio of 89 and 107, NO^+^ at m/z 118, and O_2_^+^ at m/z 88 and 106. In the case of ethanol, different product ions were selected, including H_3_O^+^ at m/z 47 and 65, and NO^+^ at m/z 45 and 63. Since the production rates of volatile ethanol and ethyl acetate kept relatively constant during the 24 h incubation (Supplementary Fig. S1), the production amount of ethanol and acetic acid in air was converted to mmol kg^−1^ based on the flushing flow rate in the experimental set-up (15 L h^−1^). The total production of ethanol and acetic acid at different time points was then calculated as the sum of their amounts in the tissue, buffer, and gas bag and expressed as mmol kg^−1^.

### 2.6. Analysis of polar metabolites

The analysis of polar metabolites was performed following the method described by Terzoudis et al. (2022a) and Bekele et al. (2014). Polar metabolites were extracted from 200 mg ground sample using 1 mL of cold methanol. The extraction was carried out in a thermomixer (Eppendorf AG, Hamburg, Germany) at 70 °C for 15 min. The resulting mixture was centrifuged at 22,000 g at 4 °C for 20 min. Then, 100 µL of the supernatant along with 20 µL of either sugar internal standard (3 g phenyl-β-D glucopyranoside/L methanol) or acid internal standard (0.1 g 3-(4-hydroxyphenyl)-propionic acid/L methanol) was dried at 50 °C under a steady nitrogen flow (Stuart, sample concentrator (SBH CON/1), Bibby Scientific Limited Stone, and Staffordshire, UK). Afterward, metabolites were derivatized by methoxymation followed by silylation following two complimentary derivatization protocols. **BSTFA protocol.** For derivatizing sugar metabolites and some acid metabolites, the dried residue was mixed with 100 µL MeOX (20 mg methoxyamine hydrochloride/mL pyridine) and incubated in a thermomixer at 30 °C for 1 h. Silylation was then performed by adding 100 µL BSTFA (N,O-bis(trimethylsilyl)trifluoroacetamide with 1 % trimethylchlorosilane) followed by incubation at 45 °C for 2 h. **MTBSTFA protocol.** For derivatization of most acid metabolites, the dried residue was re-dissolved in 100 µL of 20 g L^−1^ MeOX and derivatized at 37 °C for 90 min followed by a 30 min treatment with 100 µL of MTBSTFA (N-tert-butyldimethylsilyl-N-methyltrifluoroacetamide with 1 % tert-butyldimethylchlorosilane) at 60 °C. Finally, 170 µl of the derivatized sample was transferred into glass vials containing deactivated glass inserts and centrifuged at 22,000 g for 3 min at 18 °C.

All derivatized samples were analyzed using a Trace 1300 GC coupled with a TSQ Duo MS (Thermo Scientific, Interscience, Belgium), and equipped with a HP-5 ms column (30 m length; 250 μm internal diameter; 0.25 μm film thickness, Supelco, Bellefonte, CA). A constant flow of 1 mL min^−1^ of helium was used as the carrier gas. 1 µL of derivatized sample was injected into the machine, and four different split modes and two different temperature profiles were applied.

To analyze sugar metabolites, a split ratio of 1:200 was employed for fructose and glucose, while sucrose and myo-inositol were analyzed at a 1:20 split ratio. The oven temperature was initially set at 90 °C for 2 min, followed by a sequential ramp that alternated between 50 °C min^−1^ and 10 °C min^−1^ until reaching a final temperature of 325 °C, at which it was maintained for 5 min. The total run time for this analysis was 17.3 min. Additionally, to analyze putrescine, α-ketoglutarate, 3-phosphoglycerate, glucose-6-phosphate and fructose-6-phosphate, a split ratio of 1:7 was used. For the metabolites which were derivatized by MTBSTFA, the split ratio of 1:7 and 1:60 were used. The oven temperature started at 80 °C for 2 min and then increased at a rate of 4 °C min^−1^ until it reached 280 °C, where it was kept for 10 min. The total run time was 62 min. In total, six sugars, eight organic acids, 19 amino acids and one polyamine compound were measured by GC-MS. Information on the metabolites identified by GC-MS analysis is presented in Supplementary Table S2.

For MS detection, selected ion monitoring mode (SIM) was used. The fragment (m/z values) encompassing the entire carbon backbone of each metabolite were selected, and then the mass range from 1 atomic mass unit (amu) lighter than unlabeled fragment to 3 amu heavier than fully labeled fragment was quantified (Long and Antoniewicz, 2019).

A mixture of pure standard compounds was prepared to create an in-house library with retention times and qualifier ions to identify metabolites following the same analysis protocols outlined above. The annotation of metabolites was conducted by comparing the acquired spectra with the in-house library, using Chromeleon 7.2.10 ES software (Thermo Scientific, Interscience, Belgium), and with the NIST (National Institute of Standards and Technology, USA) library (version 14).

### 2.7. Metabolites concentration and labeling enrichment calculation

The concentrations of metabolites were determined by creating seven-point calibration curves using standards for all identified compounds. In order to correct the data for the presence of naturally occurring stable isotopes and impurities of the tracer substrate, IsoCorrectoR was used (Heinrich *et al*., 2018). The percentage ^13^C enrichment was calculated from the total abundance of ^12^C and ^13^C ions in a particular metabolite pool (Araújo *et al*., 2014).

### 2.8. Statistical analysis

Statistical analysis was performed using SPSS 21.0 (SPSS Inc., Chicago, IL, USA). Tukey’s honestly significant difference (HSD) test was used to calculate significant differences between three tissues or between different O_2_ at *p* < 0.05. Significant differences between 0 h and 24 h were calculated using an independent t-test. To reveal the correlation structure of the data a principal component analysis (PCA) was conducted using Unscrambler^®^ X (version 10.5.1, CAMO A/S, Trondheim, Norway). Prior to multivariate statistical analyses, the relative responses of the metabolites were mean centered and standardized to unit variance.

## 3. Results

### 3.1. Comparison of initial concentration of identified compounds in three tissues

In order to summarize the main differences in the distribution of metabolite concentrations in the three tomato fruit tissues, a heat-plot like visualization was made for the three different tissues (Fig. 1 and Supplementary Fig. S2). Based on the statistical analysis (Supplementary Table S3), metabolites with more than 2-fold differences between three tissues are shown in Fig. 1, and the remaining metabolites are shown in Supplementary Fig. S2.

**Fig. 1.**
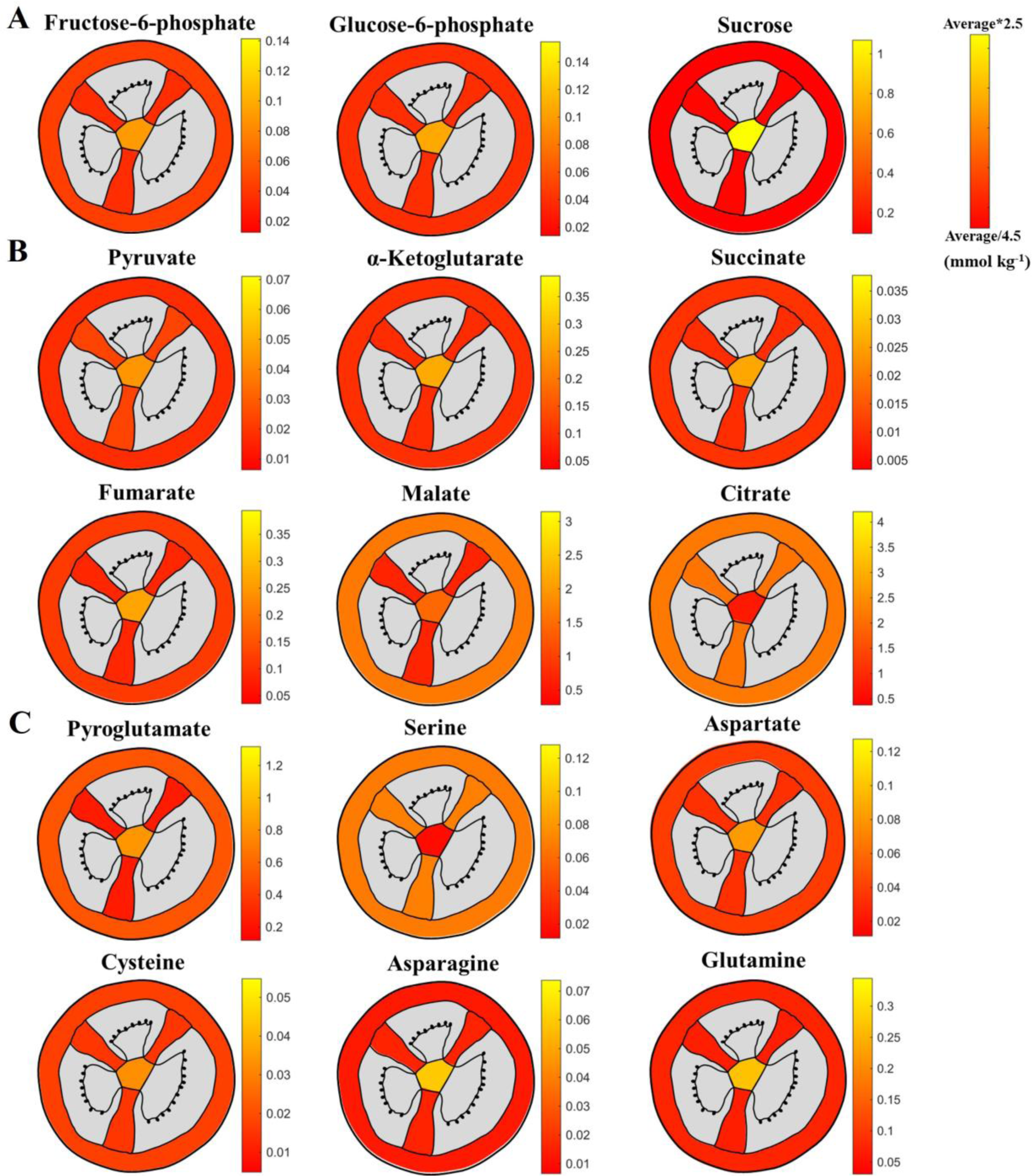
Heat plot showing the distribution of concentration for identified sugars (A), organic acids (B), and amino acids (C) with more than 2-fold difference between the columella, septa, and mesocarp tissues of tomato fruit at the beginning of the incubation period (t=0 h). The scale for each plot ranges from 1/4.5 to 2.5 times the average of the metabolite concentration in the three tissues. Tissues not measured are marked in grey.

Comparing the initial concentrations of sugars in three tissues, the concentrations of fructose-6-phosphate, glucose-6-phosphate, and sucrose were significantly higher in the columella (Fig. 1A). Notably, the concentration of sucrose in the columella was approximately 10 times higher than in the septa and mesocarp tissues (Supplementary Table S3). The concentrations of fructose, glucose and myo-inositol were only slightly different between the three tissues (Supplementary Fig. S2A, Supplementary Table S3). In terms of organic acids, the concentrations of pyruvate, α-ketoglutarate, succinate and fumarate were notably higher in columella tissue, whereas the citrate concentration was significantly lower in the columella as compared to septa and mesocarp tissues. In addition, the malate concentration was significantly lower in septa tissue (Fig. 1B, Supplementary Table S3). Most of amino acids were found in higher concentrations in columella tissue, whereas the concentrations were similar in septa and mesocarp tissues (Supplementary Table S3). Notably, the concentrations of pyroglutamate, aspartate, cysteine, asparagine, and glutamine were notably higher in the columella. However, serine concentration was significantly higher in septa and mesocarp tissues (Fig. 1C, Supplementary Table S3). As for fermentation metabolites, ethanol and acetic acid had similar concentrations in the three tissues (Supplementary Fig. S2D, Supplementary Table S3).

### 3.2. The global metabolite difference between three tissues and between different O_2_ conditions as revealed by multivariate statistical analysis

Multivariate analysis was conducted to determine latent relationships between the metabolites measured and the effect of different tissues and O_2_ conditions on their concentration profiles (Fig. 2). The amount of variation covered by PC1 and PC2 was 43 and 24 %, respectively, covering 67 % of the total variance. The PCA biplot, based on the concentrations of the identified metabolites, exhibited a clear segregation of columella tissue from septa and mesocarp tissues. Based on the projected scores for the first two PCs, the septa and mesocarp overlapped during incubation. A clear separation was observed between the different O_2_ conditions, with the 21 kPa O_2_ conditions being significantly different from those at 1 kPa and 0 kPa, while the 1 kPa and 0 kPa conditions were more similar to each other. Additionally, there was an effect of sampling time on metabolite concentrations, which was more pronounced at 1 kPa and 0 kPa than at 21 kPa.

**Fig. 2.**
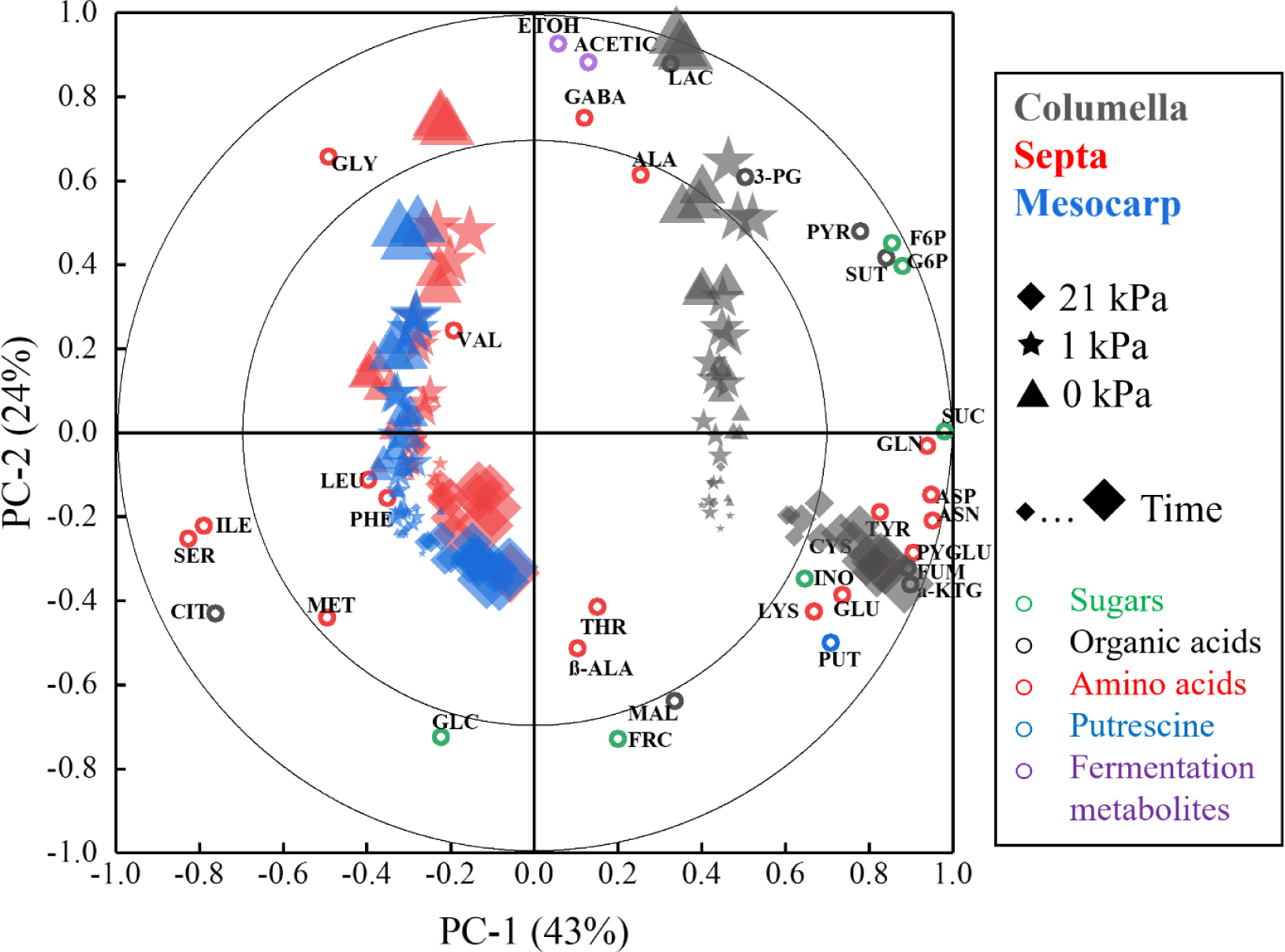
PCA biplot displaying variation in metabolite levels between columella (black), septa (red), and mesocarp (blue) tissues after incubation with ^13^C_6_ glucose for 0 h, 2 h, 4 h, 8 h, 12 h and 24 h at 21 kPa (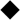), 1 kPa (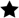) and 0 kPa (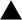) O_2_. The scores (diamonds, sparkles, and triangles) represent the different sampling points and were shown as the size of the diamonds, sparkles, and triangles increases with time, starting from 0 h to 24 h. The loading of the PCA (open circles) represents the identified metabolites (green for sugars, black for organic acids, red for amino acids, blue for putrescine, and purple for fermentation metabolites). The percentage of the explained variances is shown on the axes. Metabolites abbreviated (F6P, fructose-6-phosphate; G6P, glucose-6-phosphate; FRC, fructose; GLC, glucose; INO, myo-Inositol; SUC, sucrose; PYR, pyruvate; LAC, lactate; GABA, γ-aminobutyrate; SUT, succinate; FUM, fumarate; MAL, malate; CIT, citrate; α-ΚΤG, α-ketoglutarate; 3-PG, 3-phosphoglycerate; ALA, alanine; GLY, glycine; ß-ALA, ß-Alanine; VAL, valine; LEU, leucine; ILE, isoleucine; PYGLU, pyroglutamate; MET, methionine; SER, serine; THR, threonine; PHE, phenylalanine; ASP, aspartate; GLU, glutamate; CYS, cysteine; ASN, asparagine; LYS, lysine; GLN, glutamine; TYR, tyrosine; PUT, putrescine; ACETIC, acetic acid; ETOH, ethanol).

By overlaying metabolite loadings with sample scores, correlations between metabolites and different tissues, different O_2_ conditions and different time points can be inferred. Most of the identified metabolites were positively correlated with columella tissue, whereas citrate and some other amino acids, such as serine, glycine, methionine and isoleucine were positively correlated with septa and mesocarp tissues. On the other hand, most metabolites exhibited a positive correlation with time at 21 kPa. However, fermentation metabolites, organic acids (pyruvate, succinate, 3-phosphoglycerate and lactate), sugars (fructose-6-phosphate and glucose-6-phosphate) and amino acids (alanine, γ-aminobutyrate and glycine) were positively correlated with time at 1 kPa and 0 kPa.

### 3.3. The responses of central carbon metabolism at different O_2_ concentrations

The O_2_ consumption and CO_2_ production rates for three tissues at different O_2_ concentrations are shown in Fig. 3. All tissues show a typical Michaelis Menten behavior, as the external O_2_ concentration decreases, the rate of O_2_ consumption decreased. Similarly, the rate of CO_2_ production initially decreased, but increased as the O_2_ concentration approached 0 kPa reflecting fermentative activity.

**Fig. 3.**
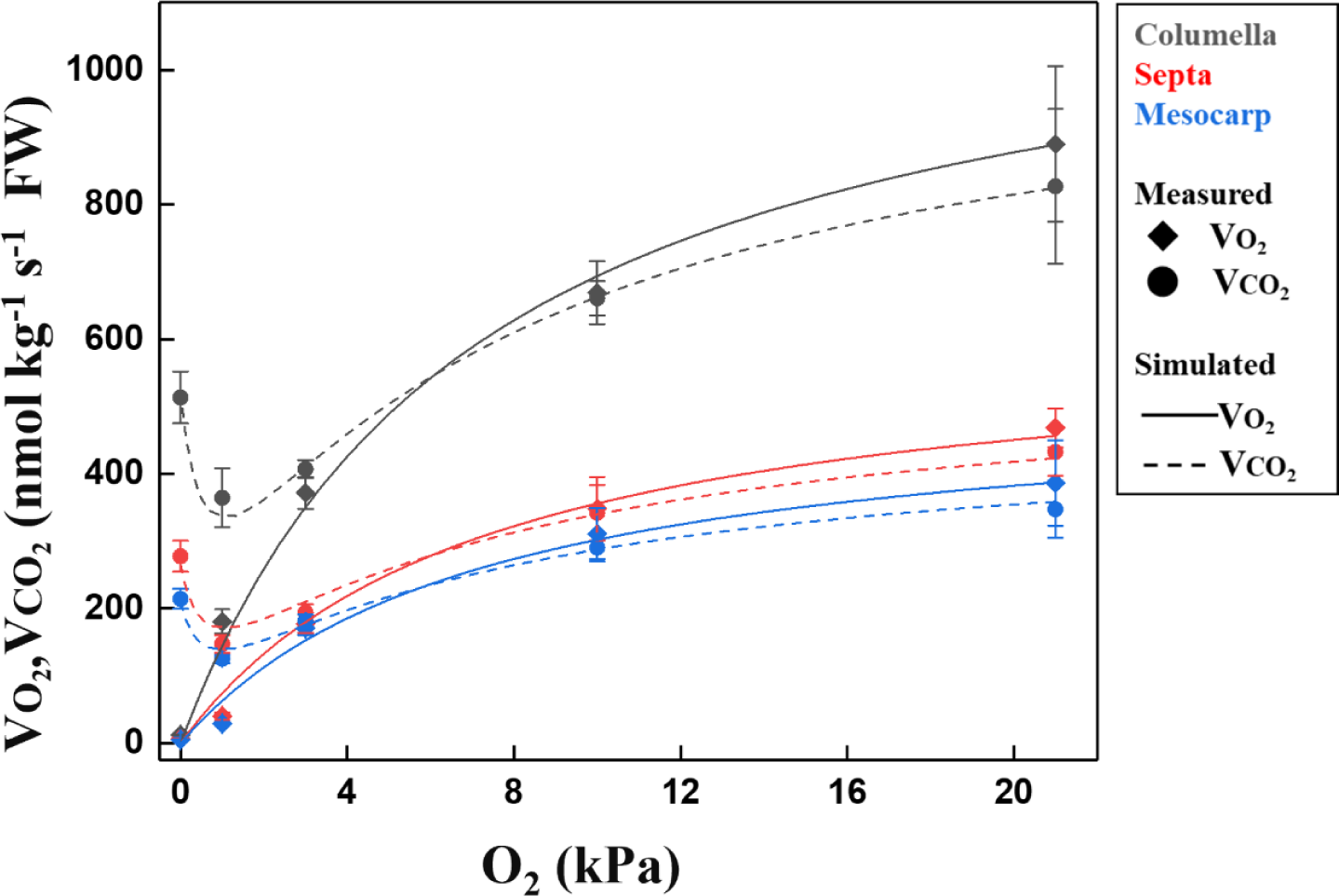
Simulated versus measured O_2_ consumption rate (*V*O_2_, diamond symbols and continuous lines) and CO_2_ production rate (*V*CO_2_, circle symbols and dotted lines) for columella (black), septa (red) and mesocarp (bule) tissues (at 18 °C) as a function of O_2_ concentration by assuming Michaelis-Menten kinetics. The symbols are measured values while the lines are simulated values. Values are means ± se of three independent measurements.

To better understand the metabolic changes in columella, septa and mesocarp tissue at 21, 1 and 0 kPa O_2_, the concentration profiles (Figs. 4A and 5, and Supplementary Figs. S3A and S4A) and label enrichment (Fig. 4B, and Supplementary Figs. S3B and S4B) of all identified metabolites at all time points were calculated. Upon incubation at 1 and 0 kPa O_2_, the metabolism responded significantly different from tissue incubated at 21 kPa O_2_.

**Fig. 4.**
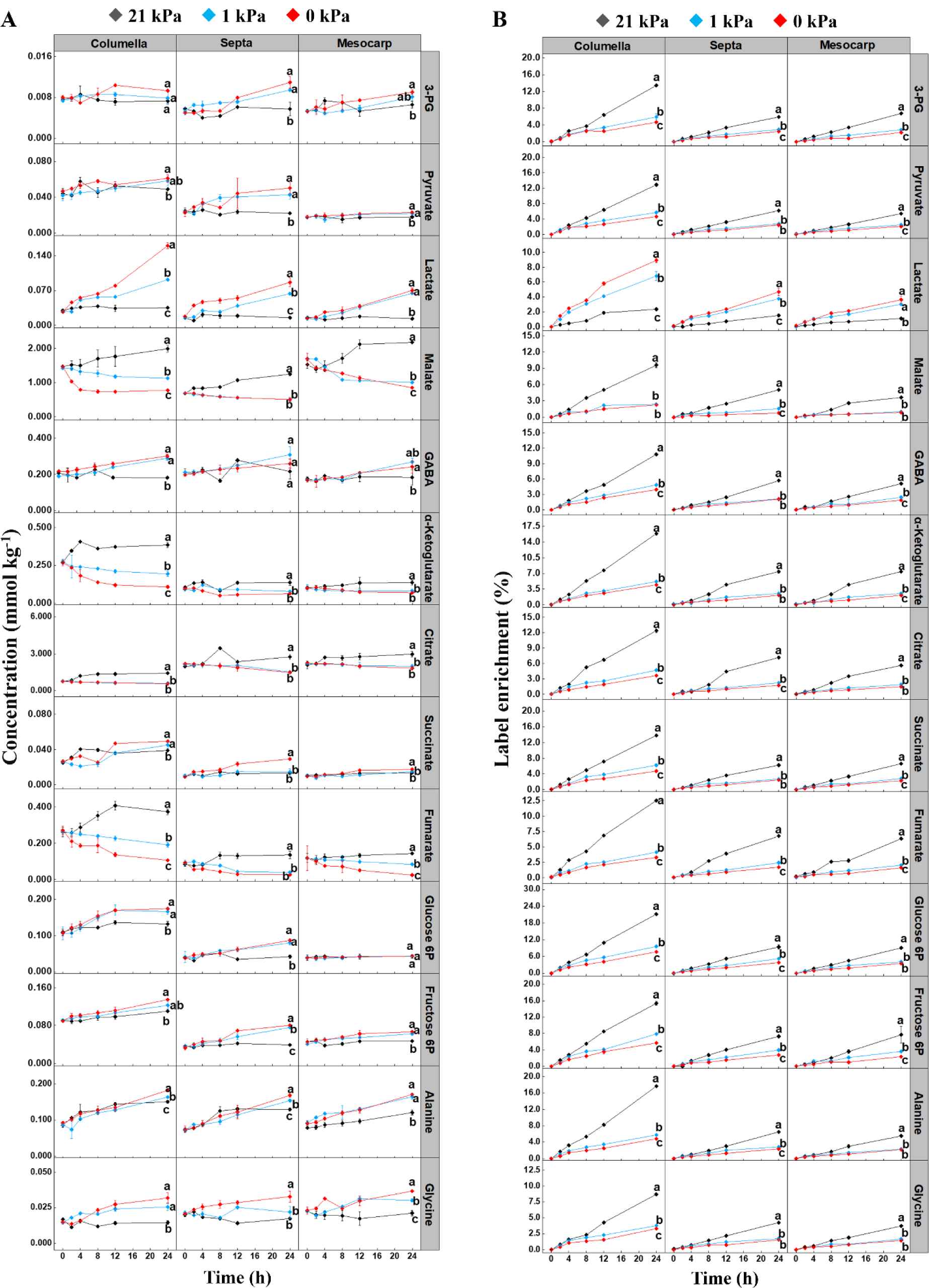
Time series showing changes in partial metabolites concentrations (A) and ^13^C label enrichment (B) in three tissues after ^13^C_6_ glucose loading at 21 kPa (black), 1 kPa (blue), and 0 kPa (red) O_2_ conditions. The temperature remained constant at 18 °C throughout the experiment. All values represent the mean of three independent replicates, and error bars indicate the standard error of the mean. Significant changes between different O_2_ concentrations in the same tissue after 24 h were based on Tukey’s honestly significant difference (HSD) test at a significant level of 0.05 and are indicated by different letters.

**Fig. 5.**
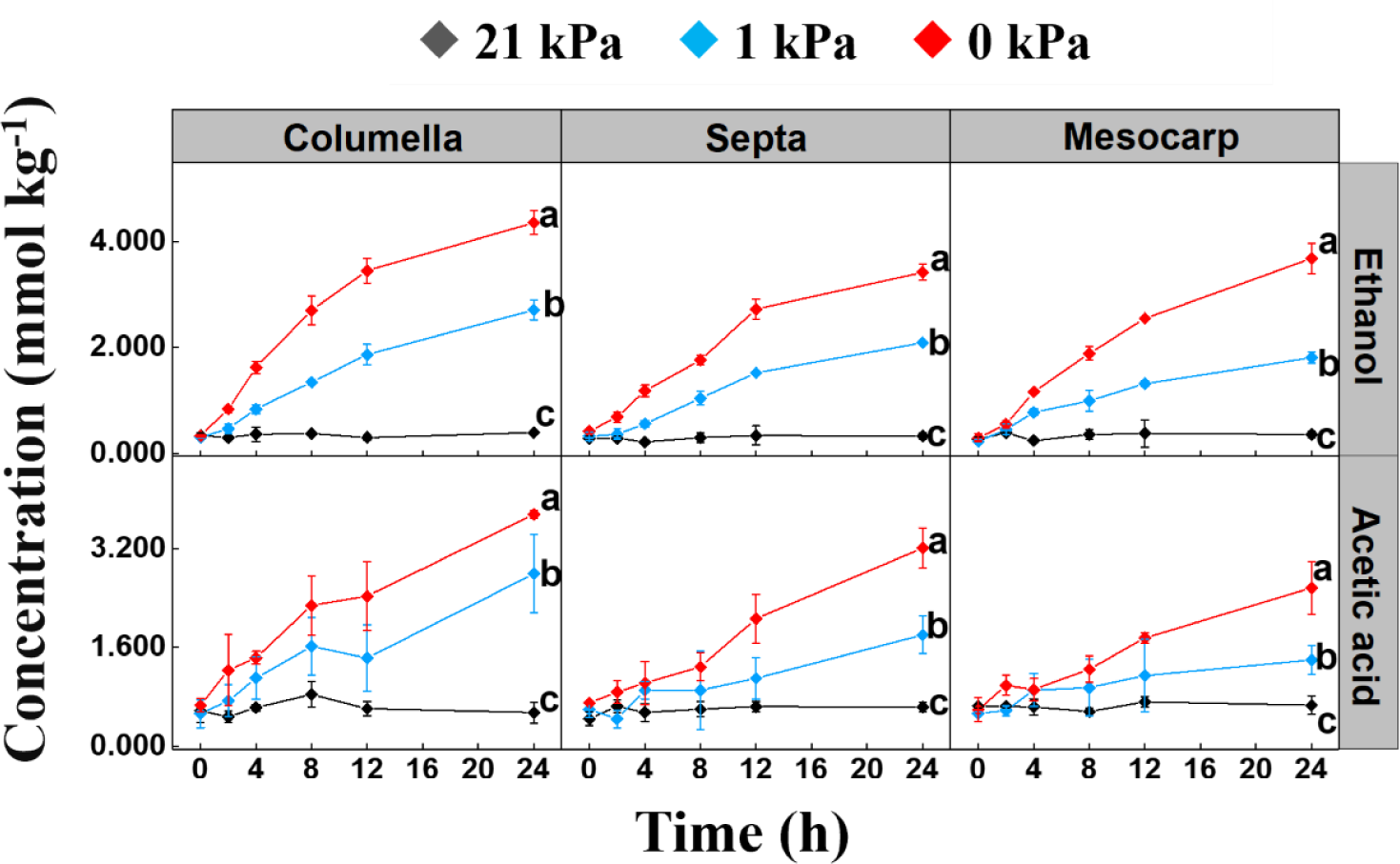
Time series showing changes in the concentration of ethanol and acetic acid in three tissues after ^13^C_6_ glucose loading at 21 kPa (black), 1 kPa (blue), and 0 kPa (red) O_2_ conditions. The temperature remained constant at 18 °C throughout the experiment. All values represent the means of three independent replicates, and error bars indicate the standard error of the mean. Significant changes between different O_2_ concentrations in the same tissue after 24 h were based on Tukey’s honestly significant difference (HSD) test at a significant level of 0.05 and are indicated by different letters.

In the three tissues, the concentration of most identified metabolites remained relatively stable over time during incubation at 21 kPa O_2_, with slight variability observed. On the contrary, there was a decreasing trend in metabolite concentrations over time at 1 and 0 kPa O_2_. Consequently, after 24 h of incubation, metabolite concentrations were generally higher under 21 kPa O_2_ conditions (Figs. 4A, 5 and Supplementary Figs. S3A and S4A). However, for sugars, the concentrations of fructose-6-phosphate and glucose-6-phosphate tended to increase over time at 1 and 0 kPa O_2_, resulting in higher concentrations after 24 h of incubation under these conditions (Fig. 4A). Similarly, among the organic acids, the concentrations of 3-PG (3-phosphoglycerate), pyruvate, lactate and succinate increased with time after 24 h incubation at 1 and 0 kPa, leading to higher concentrations at 1 and 0 kPa (Fig. 4A). As for amino acids, GABA (γ-aminobutyrate), alanine, glycine and valine concentrations also showed an increasing trend with time at 1 and 0 kPa, resulting in higher levels after 24 h incubation at these conditions (Fig. 4A and Supplementary Fig. S4). Regarding fermentation metabolites, both ethanol and acetic acid concentrations showed a similar increasing trend over time at 1 and 0 kPa, leading to significantly higher concentrations in 1 and 0 kPa after 24 h incubation (Fig. 5).

In terms of fractional enrichment, all identified compounds exhibited enrichment under 21, 1 and 0 kPa O_2_ across all three tissues. As the concentrations of ethanol and acetic acid were determined using enzymatic kits no information on the labeling enrichment of these compounds was available. The label enrichment of almost all metabolites increased over time in the three tissues under three O_2_ conditions, being significantly higher under 21 kPa O_2_ (Fig. 4B and Supplementary Figs. S3B and S4B). However, the ^13^C label enrichment of lactate was significantly higher in low O_2_ conditions (1 and 0 kPa) as compared to 21 kPa O_2_ (Fig. 4B).

After 24 h of incubation, total exogenous ^13^C_6_ glucose uptake equivalents were calculated for three tissues by summing the labeled carbon concentrations of all metabolites and converting them to ^13^C_6_ glucose concentrations. To this end, the ^13^C label enrichment of ethanol and acetic acid was assumed to be equivalent to the ^13^C label enrichment of pyruvate which is their precursor.

Significantly higher uptake of ^13^C_6_ glucose was observed under 21 kPa O_2_ conditions (Fig. 6). This lower uptake of ^13^C_6_ glucose under low O_2_ conditions results by definition in a reduction of ^13^C labeling of all downstream metabolites. For this reason, the ^13^C uptake efficiency of labeled metabolite after 24 h of incubation was calculated for each of the three tissues by multiplying the metabolite concentration at 24 h by the fractional enrichment and dividing by total exogenous ^13^C_6_ glucose uptake equivalents. For most metabolites in all tissues the ^13^C uptake efficiency was the highest at 21 kPa O_2_ (Fig. 7 and Supplementary Fig. S5). However, for sugar phosphates (fructose 6-phosphate and glucose 6-phosphate), some organic acids (3-PG, pyruvate, lactate and succinate), and amino acids (GABA, alanine and glycine) (Fig. 7), the ^13^C uptake efficiency was higher at 1 and 0 kPa O_2_.

**Fig. 6.**
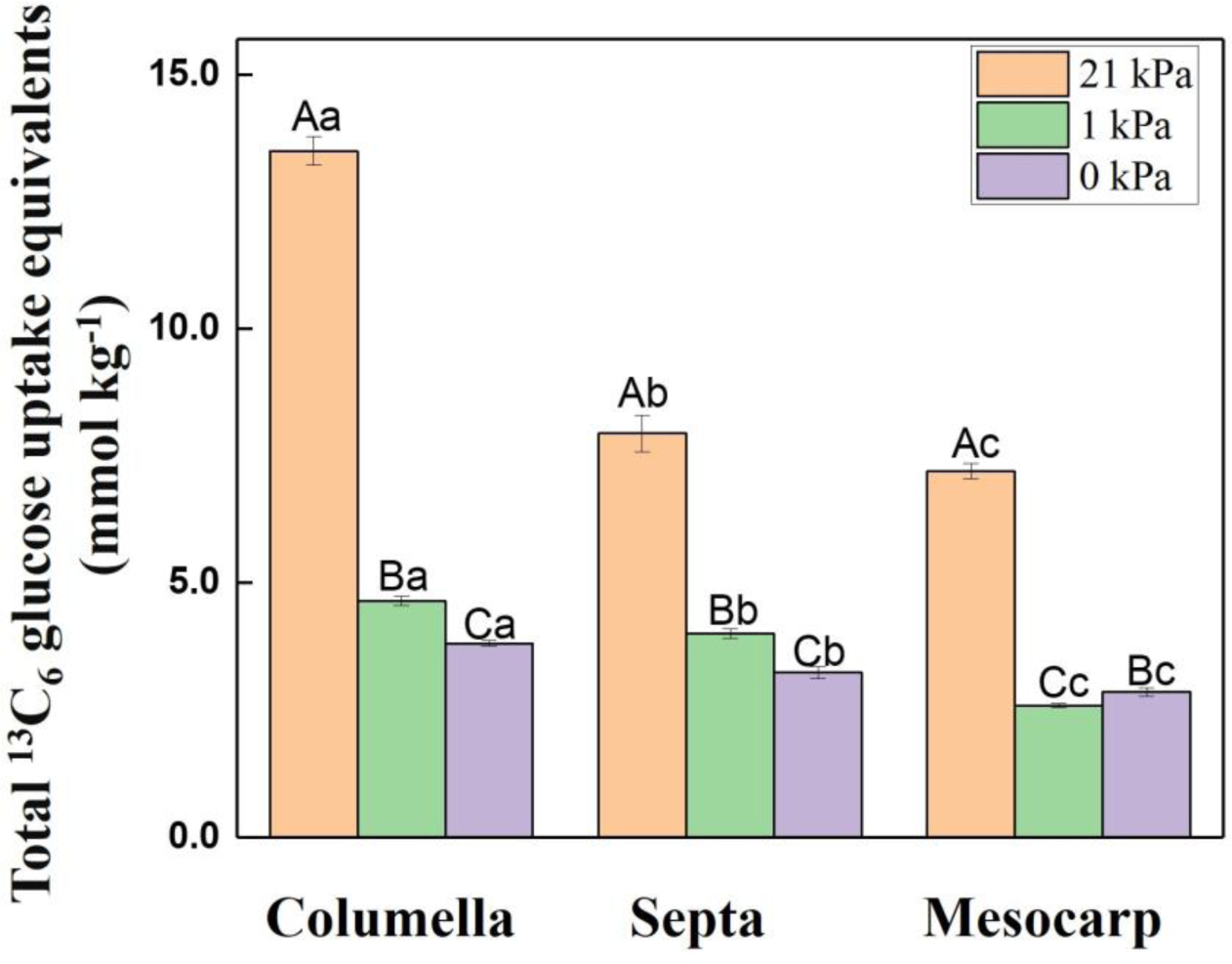
Total ^13^C_6_ glucose uptake equivalents by three tissues after 24 h incubation under three O_2_ conditions. The uptake of ^13^C_6_ glucose values were calculated for three tissues by summing the labeled carbon concentrations of all metabolites and converting them to ^13^C_6_ glucose concentrations. Error bars indicate the standard error of the mean. Significant changes between different O_2_ concentrations and tissues after 24 h were based on Tukey’s honestly significant difference (HSD) test at a significant level of 0.05. Different capital letters show significant differences between three O_2_ conditions for the same tissue, and different lower letters indicate significant differences between three tissues in the same O_2_ condition.

**Fig. 7.**
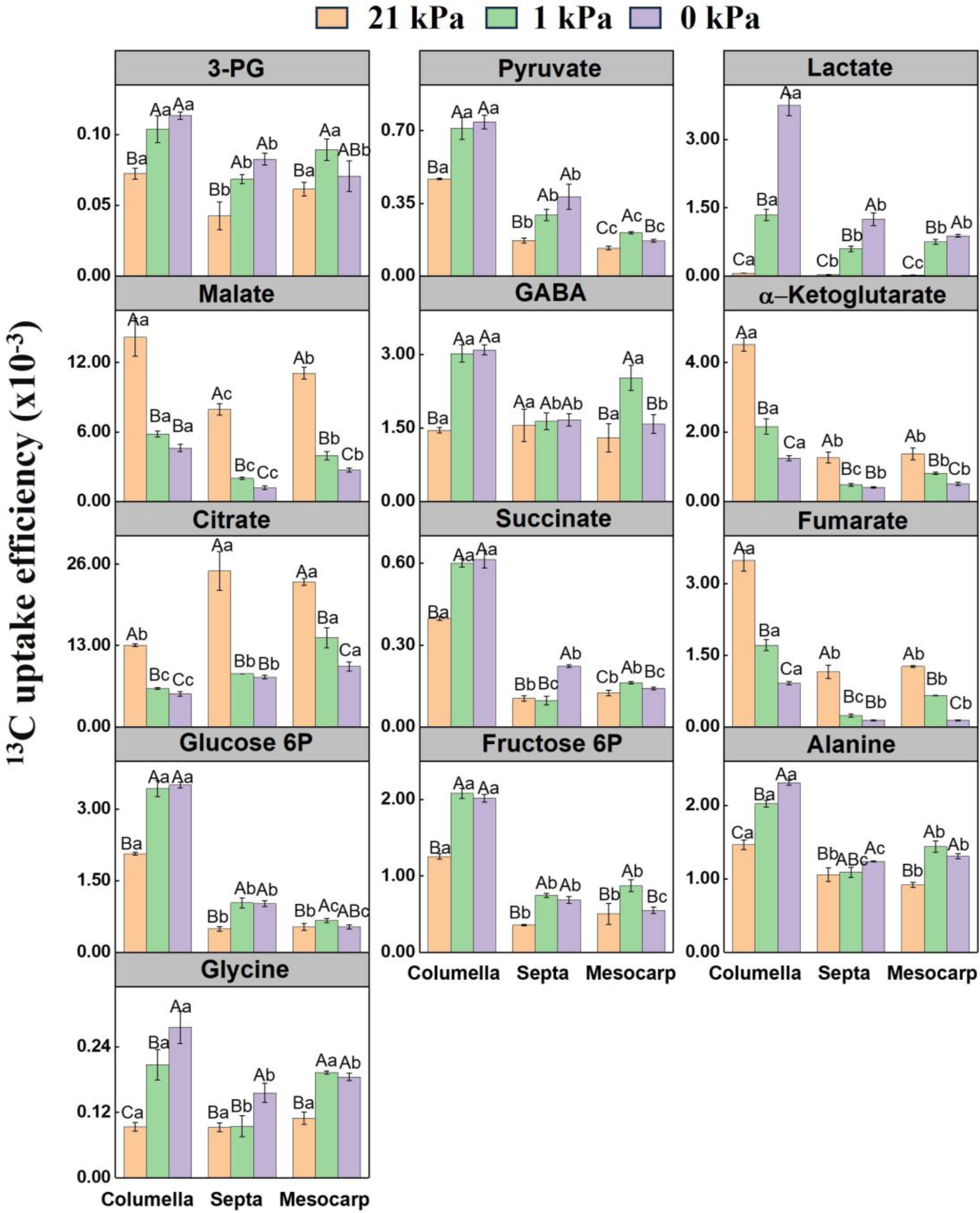
^13^C uptake efficiency of partial metabolites in three tissues after 24 h of incubation under three O_2_ conditions. The ^13^C uptake efficiency was calculated by multiplying the metabolite concentration by the enrichment and dividing by the total ^13^C_6_ glucose uptake equivalents. Error bars indicate the standard error of the mean. Significant changes between different O_2_ concentrations and tissues were based on Tukey’s honestly significant difference (HSD) test at a significant level of 0.05. Different capital letters show significant differences between three O_2_ conditions for the same tissue, and different lower letters indicate significant differences between three tissues in the same O_2_ condition.

### 3.4. The differences of the central carbon metabolism in three tissues

The respiratory parameters of *V*_max,O2_, *V*_max,CO2_, *K*_m,O2_, *RQ* and *K*_m,f,O2_ were obtained using least square nonlinear optimization using OptiPa (Hertog *et al*., 2007). The estimated Michaelis-Menten parameters are shown in Supplementary Table S4. The columella showed higher values of *V*_max,O2_ and *V*_max,CO2_ than septa and mesocarp tissues.

*K*_m,O2_, *RQ* and *K*_m,f,O2_ were the same for the three tissues.

The statistical analysis (Supplementary Table S5) showed that at 21 kPa O_2_, after 24 h of incubation, the concentration of glucose-6-phosphate, fructose-6-phosphate, myo-inositol, tyrosine, leucine, methionine, and cysteine changed more significantly in columella tissue. Meanwhile, threonine concentration changed more significantly in the septa, and 3-phosphoglycerate concentration changed significantly in the mesocarp tissue. In addition, the concentration of ß-alanine, serine and tyrosine increased significantly in both columella and septa tissues. Under low O_2_ conditions, more of the identified compounds exhibited significant concentration changes in columella and septa tissues as compared to mesocarp tissue. Regarding the fractional enrichment of all identified compounds, the statistical analysis (Supplementary Tables S6-8) revealed significantly higher levels in columella tissue than in septa and mesocarp tissues. The ^13^C label of most metabolites in the septa tissue was similar to that of mesocarp tissue after 24 h of incubation.

Total exogenous ^13^C_6_ glucose uptake equivalents were the highest in columella tissue, followed by the septa, and the lowest in mesocarp tissue (Fig. 6). The ^13^C uptake efficiency of most metabolites was higher in columella tissue, except for β-alanine, leucine, isoleucine, methionine, serine, glutamate and citrate, whereas the ^13^C uptake efficiency of most metabolites was similar in septa and mesocarp tissues (Fig. 7 and Supplementary Fig. S5).

### 3.5. An integrated overview of the tissue-specific metabolic response to low O_2_ stress

A metabolic network heat map was constructed using the initial and end metabolite levels and the end ^13^C label accumulation throughout the experiment, to give a general overview of the main metabolic difference between three tomato tissues under 21 kPa (Fig. 8A), 1 kPa (Fig. 8B) and 0 kPa (Fig. 8C) O_2_. The metabolic scheme covers the main central carbon metabolic pathways positioning all the metabolites quantified in this study.

**Fig. 8.**
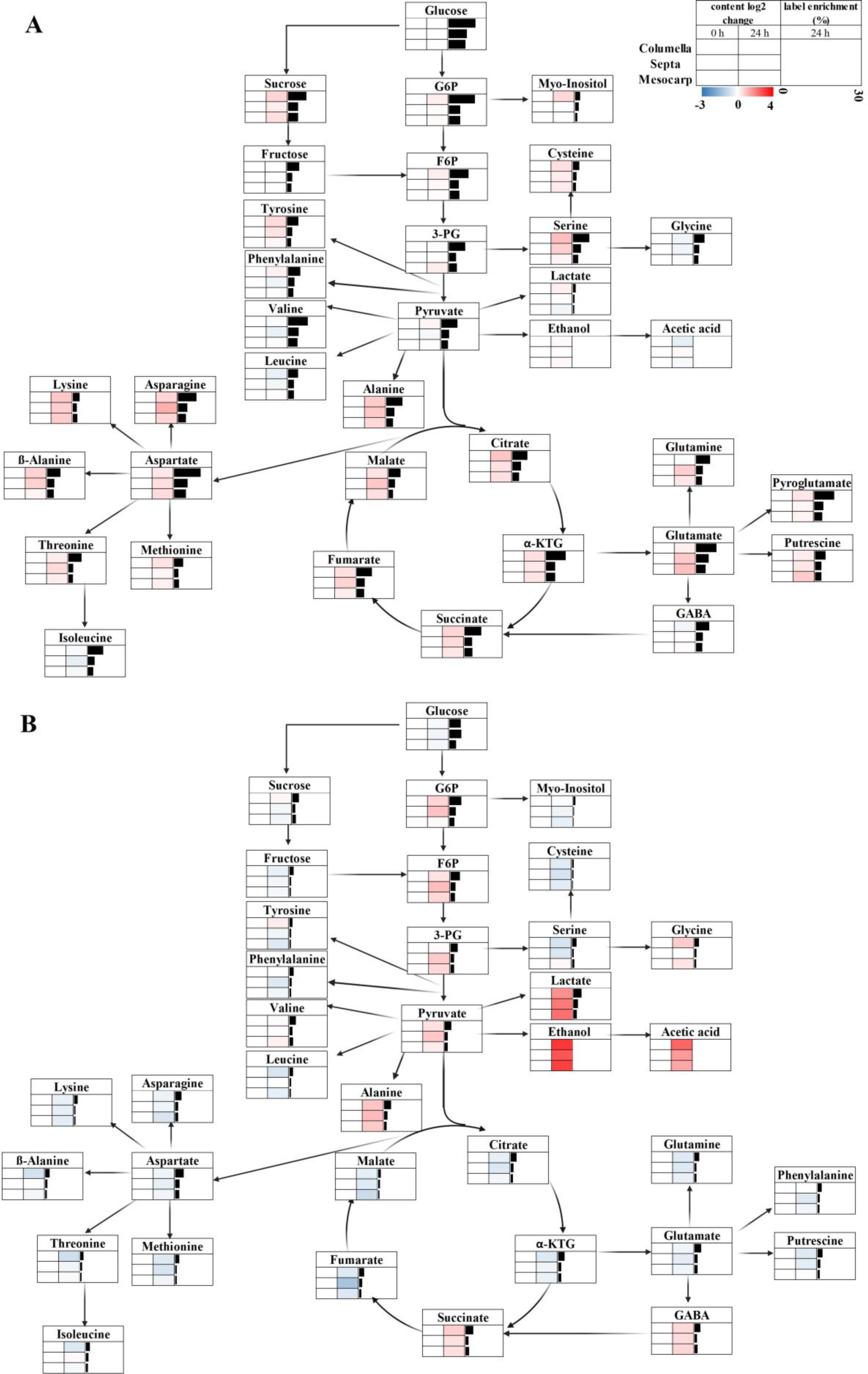
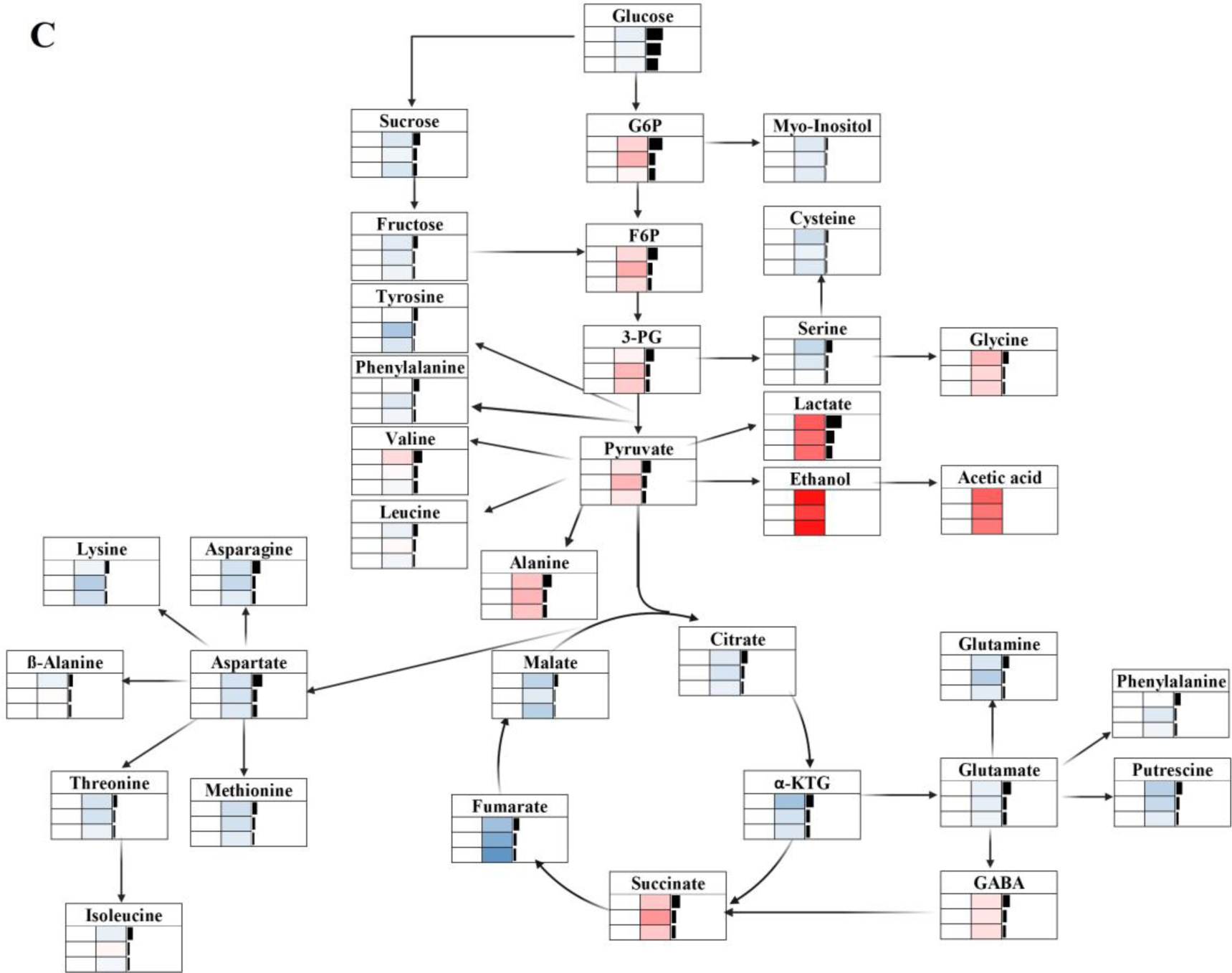
Heat map scheme of the simplified central carbon metabolism in tomato tissue showing changes in concentration and ^13^C enrichment of central carbon metabolites after 24 h incubation under 21 kPa (A), 1 kPa (B), and 0 kPa (C) O_2_. Heat maps represent the log2 fold change in the concentration of the measured metabolites as compared to the initial concentration in each tissue. The heatmap boxes show the metabolite concentration in columella, septa, and mesocarp tissue at 0 h and 24 h. The color scale (blue to red) was applied to heat map sets indicating metabolite concentration. The bar chart represents the mean of fractional enrichment of all labeled metabolites in columella, septa, and mesocarp tissue after incubation with ^13^C_6_ glucose for 24 h. The labeling levels were all expressed in a range from 0–30 %. Concentrations of ethanol and acetic acid were measured using a spectrophotometric kit, no information on labeling enrichment was available. Metabolites abbreviated (F6P, fructose-6-phosphate; G6P, glucose-6-phosphate; 3-PG, 3-phosphoglycerate; GABA, γ-aminobutyrate; α-ΚΤG, α-ketoglutarate).

At 21 kPa O_2_, the highest labeled fractions of metabolites, with the exception of lactate, and the smallest fold changes in concentration were observed for all three tissues (Fig. 8A). Under 0 and 1 kPa O_2_, the accumulation of glycolytic and fermentative metabolites increased. In addition, the accumulation of the glycolysis-derived amino acids alanine and glycine increased. In contrast, a depletion was observed of most of the intermediates from the TCA cycle and their associated amino acids. Only succinate showed a clear accumulation which was linked to an accumulation of GABA. Overall, labeling of the various metabolites was reduced, with milder effects observed in the upper part of glycolysis. Notably, there was an increased enrichment of labeling for lactate (Figs. 8B and 8C). Furthermore, in all three tissues, the columella exhibited a higher fraction of labeled metabolites as compared to septa and mesocarp tissues.

## 4. Discussion

Integrating metabolome measurements with isotopic labeling data enables the interpretation of metabolic responses through alterations in relevant pathway fluxes (Ampofo-Asiama *et al*., 2014; Heux *et al*., 2017). In this study, the combination of metabolomics and isotopic labeling data was employed to explore possible tissue differentiation with regard to the central carbon metabolism and study the impact of low O_2_ stress on the fluxes through the central carbon metabolism.

### 4.1. Low O_2_ stress induces glycolysis upregulation and initiates fermentative

In low O_2_ conditions, the activity of the mETC slows down due to the lack of O_2_ as the final electron acceptor, resulting in decreased production of ATP and NAD^+^. To avoid an energy crisis, organisms shift from aerobic metabolism to anaerobic metabolism (fermentation) which mainly produces ATP through glycolysis and regenerates NAD^+^ through fermentation (Bailey-Serres and Voesenek, 2008; Gupta *et al*., 2009; Jethva *et al*., 2022).

In this study, the accumulation of glycolysis intermediates (fructose-6-phosphate, glucose-6-phosphate, 3-phosphoglycerate and pyruvate) at 1 and 0 kPa suggests inhibition of downstream pathways following glycolysis (Fig. 4A). Similar accumulations of glycolytic intermediates under low O_2_ stress have been noted in other plant systems (Oms-Oliu *et al*., 2011; Loreti and Perata, 2020; Terzoudis *et al*., 2022b,a). However, the increased concentration of glycolytic intermediates at 1 and 0 kPa O_2_, coupled with an absence of a corresponding increase in ^13^C-label, indicate a reduced uptake rate of labeled substrate by the cells and an increased release from unlabeled intracellular sources (Fig. 4B). This observation is further supported by the increase in sucrose concentrations at 21 kPa, indicating the storage of glucose absorbed from the medium. Conversely, at 1 and 0 kPa, there is a notable decrease in sucrose levels, suggesting that under these conditions sucrose is broken down to provide substrate for subsequent reactions (Fig. S3A and Supplementary Table S5). Under energy-limited conditions, this breakdown is mainly catalyzed by sucrose synthase, which is responsible for degrading sucrose (Santaniello *et al*., 2014). Some studies have shown that the genes encoding sucrose synthases are upregulated under low O_2_ conditions in various plant species, including *Arabidopsis* (Baud *et al*., 2004), potato (Bologa *et al*., 2003) and rice (Guglielminetti *et al*., 1995). In addition, calculations of the total ^13^C_6_ glucose uptake equivalents after 24 h of incubation also confirmed the decrease of ^13^C_6_ glucose uptake under low O_2_ conditions (Fig. 6). Under low O_2_ conditions, organisms have to reduce their energy consumption to ensure survival (Geigenberger, 2003; Plaxton and Podestá, 2006; Gupta *et al*., 2009). However, the uptake of ^13^C_6_ glucose is an energy-dependent, carrier-mediated process (Brown *et al*., 1997). Therefore, the cells will take up less exogenous ^13^C_6_ glucose and utilize more preexisting intracellular glucose to reduce the consumption of energy under low O_2_ conditions. Similarly, ^13^C_6_ glucose uptake in tomato leave cells and ^14^C sucrose uptake in potato tubers were reduced under low O_2_ conditions (Geigenberger *et al*., 2000; Ampofo-Asiama *et al*., 2014). Although the lower ^13^C_6_ glucose uptake reduced the ^13^C-label in each metabolite, the ^13^C uptake efficiency in fructose-6-phosphate, glucose-6-phosphate, 3-phosphoglycerate and pyruvate was higher under low O_2_ conditions (Fig. 7). This result further confirms the upregulation of glycolysis under low O_2_ stress. Furthermore, although the accumulation of glycolysis intermediates, such as 3-phosphoglycerate, was not always significantly higher under 1 and 0 kPa O₂ conditions compared to 21 kPa (Fig. 4A), the significantly higher ¹³C uptake efficiency of these metabolites (Fig. 7) still indicates an upregulation of glycolysis under low O₂ stress. This finding emphasizes the importance of using isotope-labeled substrates in plant metabolic research, as the labeling information reveals dynamic changes that may not become obvious from the metabolite concentrations alone.

The increase of lactate, ethanol and acetic acid concentration under 1 and 0 kPa O_2_ indicates the initiation of fermentation metabolism (Figs. 4A and 5). In response to low O_2_ conditions, to counteract the buildup of cytosolic pyruvate and maintain continuous NAD^+^ regeneration, lactate dehydrogenase (LDH), alcohol dehydrogenase (ADH), and pyruvate decarboxylase (PDC) were activated and lactate, ethanol and acetaldehyde accumulated (Tadege *et al*., 1999; Jethva *et al*., 2022). This result has been well observed in apple, pear and peppers during post-harvest storage in low O_2_ conditions (Imahori *et al*., 2002; Brizzolara *et al*., 2017; Saquet and Farroupilha, 2017). In addition, ^13^C-label enrichment of lactate increased under low O_2_ conditions indicating the upregulation of LDH (Fig. 4B). Similar responses were observed in apple and tomato cell cultures (Ampofo-Asiama *et al*., 2014; Terzoudis *et al*., 2022b). Under low O_2_ conditions, the primary metabolism directs carbon atoms from the pyruvate pool towards the production of lactate, ethanol and acetic acid. However, due to the distinct volatility of fermentation by-products, the amount accumulated in the tissue is not representing their total amounts produced. To this end, analyses of tissue, medium and headspace were combined to obtain their overall production. As the high salt containing medium samples are not compatible with GC-MS, the concentrations of ethanol and acetic acid were determined using enzyme kits. Combined with the headspace samples being measured using SIFT-MS, information on ^13^C label enrichment of ethanol and acetic acid is lacking. Therefore, while approximating total carbon uptake, it was assumed that the ^13^C label enrichment of ethanol and acetic acid was comparable to that of pyruvate. There is limited information on the broad metabolic response of intact tomato fruit to low O_2_ stress. However, studies have observed increased glycolytic activity and initiation of fermentation in whole tomato fruit, reflected in both metabolic levels and gene expression (Imahori *et al*., 2003; Pegoraro *et al*., 2012), which is consistent with the responses observed at the tissue level in this study.

### 4.2. Low O_2_ stress stimulates alanine and glycine synthesis

The accumulation of lactate typically leads to a decrease in cytosolic pH, and ethanol fermentation consumes carbon to produce metabolically inactive end products (Roberts *et al*., 1984). Previous studies have found that changes in amino acid metabolism can reduce these negative effects of fermentative metabolism. Specifically, alanine metabolism plays a crucial role in maintaining glycolytic flux. This process involves the conversion of pyruvate to alanine, facilitated by the enzyme alanine aminotransferase (AlaAT) (Vandendriessche *et al*., 2013; Cukrov *et al*., 2016; Brizzolara *et al*., 2017; Boeckx *et al*., 2019). Glutamate serves as the amine group donor in this reaction (Rocha *et al*., 2010a). In this study, the drop in concentration of glutamate (Supplementary Fig. S4A) in low O_2_ conditions coincided with the increase in the concentration of alanine (Fig. 4). The accumulation of alanine under low O_2_ conditions suggests an upregulation of alanine metabolism in this study. In addition, the higher ^13^C uptake efficiency of alanine under 1 and 0 kPa O_2_ also supported the upregulation of AlaAT (Fig. 7). Similar alanine accumulation has been observed in many plant species under low O_2_ conditions (Miyashita *et al*., 2007; Limami *et al*., 2008; Rocha *et al*., 2010b,a; De Ollas *et al*., 2021).

Additionally, a significant accumulation of glycine was observed at 1 and 0 kPa in this study (Fig. 4) together with an increased ^13^C uptake efficiency (Fig. 7). The accumulation of glycine has also been reported in molecular responses of rice and wheat coleoptiles to anoxia (Shingaki-Wells *et al*., 2011). However, the mechanism behind glycine accumulation under hypoxic conditions remains unclear based on the available studies.

### 4.3. Low O_2_ stress decreases TCA cycle activity and upregulates GABA shunt, leading to succinate accumulation

Moreover, a consequence of low O_2_ stress is the reduced TCA cycle activity, which results in an insufficient supply of substrates for ATP production by the mETC, and a decrease in NAD^+^ production by the mETC, which in turn limits the activity of the TCA cycle (Boeckx *et al*., 2019). This decreased TCA cycle activity has been widely observed across various plant species subjected to low O_2_ stress, such as tomato, pear and lotus japonicus (Rocha *et al*., 2010a; Ampofo-Asiama *et al*., 2014; Terzoudis *et al*., 2022a; Mahomud *et al*., 2024). In this study, the malate, citrate, α-ketoglutarate and fumarate concentration decreased in response to low O_2_ stress, along with low ^13^C label enrichment and the lower ^13^C uptake efficiency (Figs. 4 and 7). These findings collectively suggest a reduction in TCA cycle activity. However, unlike other intermediates of the TCA cycle, the succinate concentration increased under low O_2_ conditions (Fig. 4A). In addition, the ^13^C uptake efficiency of succinate was higher under low O_2_ conditions (Fig. 7). Accumulation of succinate appears to be a general phenomenon in plants exposed to low O_2_ stress, as similar observations have been found in many other plant species (Rocha *et al*., 2010a; António *et al*., 2016; Cukrov *et al*., 2016; Boeckx, 2018). Accumulation of succinate under hypoxic conditions may be due to disruption of the cyclicity of the TCA cycle or upregulation of the GABA shunt (Terzoudis *et al*., 2022b).

Under low O_2_ stress, the TCA cycle no longer operates in a circular way as suggested by Sweetlove et al. (2010) but is split into an oxidative and a reductive part both facilitating the production of succinate. Terzoudis et al. (2022b) showed that in ‘Conference’ pear subjected to 24 h period under 0.2 kPa O_2_, malate and fumarate remained unlabeled. This observation suggests fumarase and malate dehydrogenase (MDH) are downregulated, leading to inactivation of the TCA cycle. In contrast, α-ketoglutarate and succinate remained constantly enriched, suggesting a sustained upregulation of isocitrate dehydrogenase (IDH), α-ketoglutarate dehydrogenase (OGDH) and succinate dehydrogenase (SDH). Disruption of the cyclic nature of TCA has been documented in a variety of organisms such as barley seeds, soya beans and Jonagold apples (Grafahrend-Belau *et al*., 2009; António *et al*., 2016; Boeckx, 2018). However, due to the lower ^13^C uptake efficiency of malate, citrate, α-ketoglutarate and fumarate, and their decreased ^13^C enrichment as well as their reduced concentration (Figs. 4 and 7), the accumulation of succinate cannot be conclusively attributed to the bifurcation of the TCA cycle.

The link between GABA and the TCA cycle via the GABA shunt pathway has been revealed in tomato and potato plants (Studart-Guimarães *et al*., 2007; António *et al*., 2016). This pathway bypasses succinyl-CoA ligase (SCS) and α-ketoglutarate dehydrogenase (OGDH). Instead, α-ketoglutarate is converted to GABA by the enzyme glutamate decarboxylase (GAD), a proton consuming reaction that stabilizes cytosolic pH (Cukrov *et al*., 2016; Brizzolara *et al*., 2017). The increased concentration and the ^13^C uptake efficiency of GABA (Figs. 4A and 7) indicates upregulation of GAD. However, under low O_2_ conditions, the reduced activity of TCA cycle limits α-ketoglutarate production, and the transamination between α-ketoglutarate and glutamate as well as the upregulation of GABA shunt which resulted in a drop in the concentration of α-ketoglutarate (Fig. 4A) and glutamate (Supplementary Fig. S4A). Subsequently, GABA undergoes transamination to form succinic semialdehyde, catalyzed by GABA transaminase (GABA-T). Finally, succinic semialdehyde is oxidized to succinate by succinate semialdehyde dehydrogenase (SSADH). In conditions of O_2_ limitation, as described in studies by Rocha et al. (2010) and Boeckx et al. (2019), the TCA cycle activity is reduced. Consequently, succinate is consumed less and thus accumulated, which is consistent with the observed increase in ^13^C uptake efficiency and accumulation of succinate in this study (Figs. 4A and 7).

### 4.4. Columella dominance: initial metabolite abundance, higher central carbon metabolism activity, and higher sensitivity to low O_2_ stress

Within this study, at the start of the experiment (*t* = 0 h), the columella tissue displayed a higher concentration of some sugars, organic acids, amino acids, and putrescine in contrast to both septa and mesocarp tissues. However, more similarities in metabolite concentrations were observed between the septa and mesocarp tissues (Fig. 1, Supplementary Fig. S2 and Supplementary Table S3). This finding is consistent with previous findings suggesting that different transcriptomic and secondary metabolomic profiles exist between columella and pericarp tissues (Moco *et al*., 2007; Shinozaki *et al*., 2018). In addition, the higher osmolality observed in columella tissue and similar osmolality observed in septa and mesocarp tissues coincide well with the different distribution of metabolite concentration in three tissues (Supplementary Table S1). It is noteworthy that the sucrose concentration in columella tissue was approximately ten times larger than in the septa and mesocarp tissues (Fig. 1 and Supplementary Table S3). This pattern of significantly higher sucrose concentration in columella tissue than in other fleshy tissues was confirmed in ‘Ailsa Craig’ tomato fruit by other authors (Chamley *et al*., 2019). The transcriptome profiles of various tomato fruit tissues further support this observation, as genes related to carbohydrate transport were found to have higher expression levels in the columella (Shinozaki *et al*., 2018). Sucrose, the major photoassimilate transported into fruit, is transported from vascular bundle to the fruit (Beckles *et al*., 2012). Given the highest degree of vascularization of columella tissue, it is likely that it has a higher concentration of sucrose than the other fruit tissues.

The highest maximal respiration rates (Supplementary Table S4) and total ^13^C_6_ glucose uptake equivalents (Fig. 6) observed in columella tissue suggest a higher activity of the central carbon metabolism in columella tissue. The result is further supported by the higher ^13^C uptake efficiency (Fig. 7 and Supplementary Fig. S5) of most metabolites and more significant changes in metabolite concentrations in the columella (Supplementary Table S5). Higher respiration rates in columella tissue than in septa and mesocarp tissues were also found for ‘Merlice’ tomato fruit (Xiao *et al*., 2024).

The columella hosts the vascular bundles, facilitating the connection between the fruit sink and the source leaves, as well as linking the maternal part of the fruit to the seeds. It plays a crucial role in phloem unloading and redistributing imported carbon to other fruit tissues (Chamley *et al*., 2019). Given its central role in the physiology of fruit, columella tissue may require more energy and carbon sources to maintain its complex functions. Starch as a transitory carbohydrate reserve was higher in columella tissue (Chamley *et al*., 2019). In addition, the number of mitochondria per plant cell varies and is related primarily to the metabolic activity of the tissue. Notably, mitochondria in highly active cells, such as phloem companion cells, may occupy up to 20 % of their cytoplasmic volume (DeWitt and Sussman, 1995; Buchanan *et al*., 2015). In this study, higher concentration of most metabolites in columella tissue may provide more carbon sources. In addition, the higher central carbon metabolic activity in columella tissue is in agreement with the high energy demand needed for phloem unloading. However, the same *K*_m,O2_, *RQ* and *K*_m,f,O2_ indicate the three tissues rely on the same underlying pathways that are, however, active to a different extent.

As the activity of the central carbon metabolism is heavily influenced by the change in O_2_ concentration (Boeckx *et al*., 2019), tissues with higher metabolic activity are expected to show a more pronounced response to changes in O_2_ levels. Indeed, the more abrupt change of the O_2_ consumption rate in response to a drop in the O_2_ concentration was observed in columella tissue (Fig. 3). In addition, the metabolic response to O_2_ became evident through significant changes in metabolite concentrations (Fig. 4A, Supplementary Figs. S3A and S4A, Supplementary Table S5) and ^13^C uptake efficiency (Fig. 7 and Supplementary Fig. S5), with the largest response in columella tissue. This increased metabolic sensitivity is consistent with the high expression of defense response genes in the columella (Shinozaki *et al*., 2018). Therefore, columella tissue with the highest metabolic activity showed the highest sensitivity to changes in O_2_ concentration, indicating its increased energy demand and metabolic flux.

## 5. Conclusion

This study set out to investigate possible tissue-specific responses of the central carbon metabolism to low O_2_ stress in tomato fruit by integrating metabolomics and labeling information. The accumulation of glycolytic intermediates and fermentation compounds and the increased ^13^C uptake efficiency under low O_2_ conditions indicated that all three tomato tissues initiated fermentation and up-regulated glycolysis. In addition, the accumulation of succinate and GABA as well as the increased ^13^C uptake efficiency and depletion of other TCA cycle intermediates indicated a downregulation of the TCA cycle and an upregulation of the GABA shunt. On the other hand, the difference in respiration rates and total ^13^C_6_ glucose uptake equivalents among the three tissues indicated that columella tissue exhibited the highest metabolic activity and sensitivity to the change in O_2_ concentration, followed by septa and mesocarp tissues. The higher activity of columella tissue reflects the key role of the columella in establishing vascular connectivity and carbon redistribution within the fruit. However, the ^13^C uptake efficiency of the various metabolites not always conformed to the expected pattern. Therefore, further work should use the experimental data to develop a kinetic pathway model of the central carbon metabolic pathways to quantitatively interpret the results and further explore tissue-specific responses of the central carbon metabolism in tomato fruit.

## Supplementary data

The following supplementary data are available at JXB online.

Table S1. Osmolality of columella, septa, and mesocarp tissues.

Table S2. GC-MS parameters related to the identified metabolites.

Table S3. Initial concentrations (mmol kg^−1^) of metabolites measured in the three tomato tissues at the beginning of the incubation period (t=0 h).

Table S4. Estimated Michaelis-Menten parameters for O_2_ consumption and CO_2_ production rates.

Table S5. Independent t-test p-values comparing metabolite concentrations measured at the beginning (t=0 h) and end (t=24 h) of the incubation period for the three tissues under three O_2_ conditions.

Table S6. Enrichment fractions (%) of all labeled metabolites in the three tomato tissues at 24 h of ^13^C_6_ glucose incubation at 21 kPa.

Table S7. Enrichment fractions (%) of all labeled metabolites in the three tomato tissues at 24 h of ^13^C_6_ glucose incubation at 1 kPa.

Table S8. Enrichment fractions (%) of all labeled metabolites in the three tomato tissues at 24 h of ^13^C_6_ glucose incubation at 0 kPa.

Fig. S1. Time series showing the production rates of ethanol and acetic acid from gas samples in three tissues at three O_2_ conditions.

Fig. S2. The distribution of some of the identified sugars (A), organic acids (B), amino acids (C), fermentation metabolites (D), and putrescine (E) concentration in three tissues at the beginning of the incubation period (t=0 h).

Fig. S3. The changes in sugar and putrescine concentrations (A) and ^13^C label enrichment (B) in three tissues after ^13^C_6_ glucose loading at three O_2_ conditions.

Fig. S4. The changes in amino acid concentrations (A) and ^13^C label enrichment (B) in three tissues after ^13^C_6_ glucose loading at three O_2_ conditions.

Fig. S5. ^13^C uptake efficiency of partial metabolites in three tissues after 24 h of incubation under three O_2_ conditions.

Method. S1. Determination of the preincubation time.

## Acknowledgments

The authors are grateful to Elfie Dekempeneer for her guidance in operating the GC-MS device, Tessa Vanempten and Cecilia Cardinez for their support in SIFT-MS measurements and Hui Xiao for support in drawing heat plots of initial metabolite distributions.

## Author contributions

XL: methodology, investigation, data curation, formal analysis, writing-original draft; KT: methodology; MLATMH: funding acquisition, supervision, conceptualization, writing-review and editing; BMN: funding acquisition, resources, conceptualization, writing-review and editing.

## Data availability

The data that support the findings of this study are available from the corresponding author upon request.

## References

Aizat WM, Ismail I, Noor NM. 2018. Omics applications for systems biology. Advances in Experimental Medicine and Biology. 51–55.

Ampofo-Asiama J, Baiye VMM, Hertog MLATM, Waelkens E, Geeraerd AH, Nicolai BM. 2014. The metabolic response of cultured tomato cells to low oxygen stress. Plant Biology 16, 594–606.

António C, Päpke C, Rocha M, Diab H, Limami AM, Obata T, Fernie AR, van Dongen JT. 2016. Regulation of primary metabolism in response to low oxygen availability as revealed by carbon and nitrogen isotope redistribution. Plant Physiology 170, 43–56.

Araújo WL, Tohge T, Nunes-Nesi A, Obata T, Fernie AR. 2014. Analysis of kinetic labeling of amino acids and organic acids by GC-MS. Methods in Molecular Biology. Humana Press Inc., 107–119.

Bailey-Serres J, Voesenek LACJ. 2008. Flooding stress: Acclimations and genetic diversity. Annual Review of Plant Biology 59, 313–339.

Baud S, Vaultier MN, Rochat C. 2004. Structure and expression profile of the sucrose synthase multigene family in Arabidopsis. Journal of Experimental Botany 55, 397–409.

Beckles DM, Hong N, Stamova L, Luengwilai K. 2012. Biochemical factors contributing to tomato fruit sugar content: A review. Fruits 67, 49–64.

Bekele EA, Annaratone CEP, Hertog MLATM, Nicolai BM, Geeraerd AH. 2014. Multi-response optimization of the extraction and derivatization protocol of selected polar metabolites from apple fruit tissue for GC-MS analysis. Analytica Chimica Acta 824, 42–56.

Beshir WF, Mbong VBM, Hertog MLATM, Geeraerd AH, Van Den Ende W, Nicolaï BM. 2017. Dynamic labeling reveals temporal changes in carbon re-allocation within the central metabolism of developing apple fruit. Frontiers in Plant Science 8, 1–16.

Boeckx J. 2018. Regulation of the respiratory metabolism of apple during (dynamic) controlled atmosphere storage.

Boeckx J, Pols S, Hertog MLATM, Nicolaï BM. 2019. Regulation of the central carbon metabolism in apple fruit exposed to postharvest low-oxygen stress. Frontiers in Plant Science 10, 1–17.

Boller T, Kende H. 1980. Regulation of wound ethylene synthesis in plants. Nature 286, 259–260.

Bologa KL, Fernie AR, Leisse A, Ehlers Loureiro M, Geigenberger P. 2003. A bypass of sucrose synthase leads to low internal oxygen and impaired metabolic performance in growing potato tubers. Plant Physiology 132, 2058–2072.

Brizzolara S, Santucci C, Tenori L, Hertog M, Nicolai B, Stürz S, Zanella A, Tonutti P. 2017. A metabolomics approach to elucidate apple fruit responses to static and dynamic controlled atmosphere storage. Postharvest Biology and Technology 127, 76–87.

Brown MM, Hall JL, Ho LC. 1997. Sugar uptake by protoplasts isolated from tomato fruit tissues during various stages of fruit growth. Physiologia Plantarum 101, 533–539.

Brunetti C, George RM, Tattini M, Field K, Davey MP. 2013. Metabolomics in plant environmental physiology. Journal of Experimental Botany 64, 4011–4020.

Buchanan BB, Gruissem W, Jones RL. 2015. Biochemistry and molecular biology of plants. American Society of Plant Biologists. 611–612.

Chamley ML, Mounet F, Deborde C, Maucourt M, Jacob D, Moing A. 2019. NMR-based tissular and developmental metabolomics of tomato fruit. Metabolites 9, 1–18.

Colombié S, Beauvoit B, Nazaret C, et al. 2017. Respiration climacteric in tomato fruits elucidated by constraint-based modelling. New Phytologist 213, 1726–1739.

Cukrov D, Zermiani M, Brizzolara S, et al. 2016. Extreme hypoxic conditions induce selective molecular responses and metabolic reset in detached apple fruit. Frontiers in Plant Science 7, 1–18.

DeWitt ND, Sussman MR. 1995. Immunocytological localization of an epitope-tagged plasma membrane proton pump (H+-ATPase) in phloem companion cells. Plant Cell 7, 2053–2067.

Fernie AR, Geigenberger P, Stitt M. 2005. Flux an important, but neglected, component of functional genomics. Current Opinion in Plant Biology 8, 174–182.

Geigenberger P. 2003. Response of plant metabolism to too little oxygen. Current Opinion in Plant Biology 6, 247–256.

Geigenberger P, Fernie AR, Gibon Y, Christ M, Stitt M. 2000. Metabolic activity decreases as an adaptive response to low internal oxygen in growing potato tubers. Biological Chemistry 381, 723–740.

Grafahrend-Belau E, Schreiber F, Koschützki D, Junker BH. 2009. Flux balance analysis of barley seeds: A computational approach to study systemic properties of central metabolism. Plant Physiology 149, 585–598.

Guglielminetti L, Perata P, Alpi A. 1995. Effect of anoxia on carbohydrate metabolism in rice seedlings. Plant Physiology 108, 735–741.

Gupta KJ, Zabalza A, Van Dongen JT. 2009. Regulation of respiration when the oxygen availability changes. Physiologia Plantarum 137, 383–391.

Heinrich P, Kohler C, Ellmann L, Kuerner P, Spang R, Oefner PJ, Dettmer K. 2018. Correcting for natural isotope abundance and tracer impurity in MS-, MS/MS- and high-resolution-multiple-tracer-data from stable isotope labeling experiments with IsoCorrectoR. Scientific Reports 8, 1–10.

Hertog MLATM, Peppelenbos HW, Evelo RG, Tijskens LMM. 1998. A dynamic and generic model of gas exchange of respiring produce: The effects of oxygen, carbon dioxide and temperature. Postharvest Biology and Technology 14, 335–349.

Hertog MLATM, Verlinden BE, Lammertyn J, Nicolaï BM. 2007. OptiPa, an essential primer to develop models in the postharvest area. Computers and Electronics in Agriculture 57, 99–106.

Heux S, Bergès C, Millard P, Portais JC, Létisse F. 2017. Recent advances in high-throughput 13C-fluxomics. Current Opinion in Biotechnology 43, 104–109.

Ho QT, Hertog MLATM, Verboven P, Ambaw A, Rogge S, Verlinden BE, Nicolaï BM. 2018. Down-regulation of respiration in pear fruit depends on temperature. Journal of Experimental Botany 69, 2049–2060.

Ho QT, Verboven P, Verlinden BE, Lammertyn J, Vandewalle S, Nicolaï BM. 2008. A continuum model for metabolic gas exchange in pear fruit. PLoS Computational Biology 4, e1000023.

Ho QT, Verboven P, Verlinden BE, Schenk A, Nicolaï BM. 2013. Controlled atmosphere storage may lead to local ATP deficiency in apple. Postharvest Biology and Technology 78, 103–112.

Imahori Y, Kota M, Ueda Y, Ishimaru M. 2002. Regulation of ethanolic fermentation in bell pepper fruit under low oxygen stress. Postharvest Biology and Technology 25, 159–167.

Imahori Y, Matushita K, Kota M, Ueda Y, Ishimaru M, Chachin K. 2003. Regulation of fermentative metabolism in tomato fruit under low oxygen stress. Journal of Horticultural Science and Biotechnology 78, 386–393.

Jethva J, Schmidt RR, Sauter M, Selinski J. 2022. Try or die: dynamics of plant respiration and how to survive low oxygen conditions. Plants 11, 205.

Limami AM, Glévarec G, Ricoult C, Cliquet JB, Planchet E. 2008. Concerted modulation of alanine and glutamate metabolism in young Medicago truncatula seedlings under hypoxic stress. Journal of Experimental Botany 59, 2325–2335.

Long CP, Antoniewicz MR. 2019. High-resolution 13C metabolic flux analysis. Nature Protocols 14, 2856–2877.

Loreti E, Perata P. 2020. The many facets of hypoxia in plants. Plants 9, 1–14.

Mahomud S, Islam N, Roy J. 2024. Heliyon Effect of low oxygen stress on the metabolic responses of tomato fruit cells. Heliyon 10, e24566.

Matas AJ, Yeats TH, Buda GJ, et al. 2011. Tissue- and cell-type specific transcriptome profiling of expanding tomato fruit provides insights into metabolic and regulatory specialization and cuticle formation. Plant Cell 23, 3893–3910.

Miyashita Y, Dolferus R, Ismond KP, Good AG. 2007. Alanine aminotransferase catalyses the breakdown of alanine after hypoxia in Arabidopsis thaliana. Plant Journal 49, 1108–1121.

Moco S, Capanoglu E, Tikunov Y, Bino RJ, Boyacioglu D, Hall RD, Vervoort J, De Vos RCH. 2007. Tissue specialization at the metabolite level is perceived during the development of tomato fruit. Journal of Experimental Botany 58, 4131–4146.

De Ollas C, González-Guzmán M, Pitarch Z, Matus JT, Candela H, Rambla JL, Granell A, Gómez-Cadenas A, Arbona V. 2021. Identification of ABA-Mediated Genetic and Metabolic Responses to Soil Flooding in Tomato (Solanum lycopersicum L. Mill). Frontiers in Plant Science 12, 1–20.

Oms-Oliu G, Hertog MLATM, Van de Poel B, Ampofo-Asiama J, Geeraerd AH, Nicolai BM. 2011. Metabolic characterization of tomato fruit during preharvest development, ripening, and postharvest shelf-life. Postharvest Biology and Technology 62, 7–16.

Pabón-Mora N, Litt A. 2011. Comparative anatomical and developmental analysis of dry and fleshy fruits of Solanaceae. American Journal of Botany 98, 1415–1436.

Pegoraro C, dos Santos RS, Krüger MM, Tiecher A, da Maia LC, Rombaldi CV, de Oliveira AC. 2012. Effects of hypoxia storage on gene transcript accumulation during tomato fruit ripening. Brazilian Journal of Plant Physiology 24, 141–148.

Plaxton WC, Podestá FE. 2006. The functional organization and control of plant respiration. Critical Reviews in Plant Sciences 25, 159–198.

Van de Poel B, Vandenzavel N, Smet C, et al. 2014. Tissue specific analysis reveals a differential organization and regulation of both ethylene biosynthesis and E8 during climacteric ripening of tomato. BMC Plant Biology 14, 11.

Quinet M, Angosto T, Yuste-Lisbona FJ, Blanchard-Gros R, Bigot S, Martinez JP, Lutts S. 2019. Tomato fruit development and metabolism. Frontiers in Plant Science 10, 1–23.

Ratcliffe RG, Shachar-Hill Y. 2006. Measuring multiple fluxes through plant metabolic networks. Plant Journal 45, 490–511.

Roberts JK, Callis J, Jardetzky O, Walbot V, Freeling M. 1984. Cytoplasmic acidosis as a determinant of flooding intolerance in plants. Proceedings of the National Academy of Sciences 81, 6029–6033.

Rocha M, Licausi F, Araújo WL, Nunes-Nesi A, Sodek L, Fernie AR, van Dongen JT. 2010a. Glycolysis and the tricarboxylic acid cycle are linked by alanine aminotransferase during hypoxia induced by waterlogging of Lotus japonicus. Plant Physiology 152, 1501–1513.

Rocha M, Sodek L, Licausi F, Hameed MW, Dornelas MC, Van Dongen JT. 2010b. Analysis of alanine aminotransferase in various organs of soybean (Glycine max) and in dependence of different nitrogen fertilisers during hypoxic stress. Amino Acids 39, 1043–1053.

Santaniello A, Loreti E, Gonzali S, Novi G, Perata P. 2014. A reassessment of the role of sucrose synthase in the hypoxic sucrose-ethanol transition in Arabidopsis. Plant Cell and Environment 37, 2294–2302.

Saquet A, Farroupilha T. 2017. Aroma volatiles of ‘Conference’ pear and their changes during regular air and controlled atmosphere storage. Revista de Ciência e Inovação 1, 55–66.

Schaffer AA, Petreikov M. 1997. Sucrose-to-starch metabolism in tomato fruit undergoing transient starch accumulation. Plant Physiology 113, 739–746.

Shingaki-Wells RN, Huang S, Taylor NL, Carroll AJ, Zhou W, Harvey Miller A. 2011. Differential molecular responses of rice and wheat coleoptiles to anoxia reveal novel metabolic adaptations in amino acid metabolism for tissue tolerance. Plant Physiology 156, 1706–1724.

Shinozaki Y, Nicolas P, Fernandez-Pozo N, et al. 2018. High-resolution spatiotemporal transcriptome mapping of tomato fruit development and ripening. Nature Communications 9, 364.

Smillie RM, Hetherington SE, Davies WJ. 1999. Photosynthetic activity of the calyx, green shoulder, pericarp, and locular parenchyma of tomato fruit. Journal of Experimental Botany 50, 707–718.

Studart-Guimarães C, Fait A, Nunes-Nesi A, Carrari F, Usadel B, Fernie AR. 2007. Reduced expression of succinyl-coenzyme A ligase can be compensated for by up-regulation of the γ-aminobutyrate shunt in illuminated tomato leaves. Plant Physiology 145, 626–639.

Sweetlove LJ, Beard KFM, Nunes-Nesi A, Fernie AR, Ratcliffe RG. 2010. Not just a circle: Flux modes in the plant TCA cycle. Trends in Plant Science 15, 462–470.

Tadege M, Dupuis I, Kuhlemeier C. 1999. Ethanolic fermentation: New functions for an old pathway. Trends in Plant Science 4, 320–325.

Terzoudis K, Hertog MLATM, Nicolaï BM. 2022a. Dynamic labelling reveals central carbon metabolism responses to stepwise decreasing hypoxia and reoxygenation during postharvest in pear fruit. Postharvest Biology and Technology 186, 111816.

Terzoudis K, Kusma R, Hertog MLATM, Nicolaï BM. 2022b. Metabolic adaptation of ‘Conference’ pear to postharvest hypoxia: The impact of harvest time and hypoxic pre-treatments. Postharvest Biology and Technology 189, 111937.

Vandendriessche T, Schäfer H, Verlinden BE, Humpfer E, Hertog MLATM, Nicolaï BM. 2013. High-throughput NMR based metabolic profiling of Braeburn apple in relation to internal browning. Postharvest Biology and Technology 80, 18–24.

Wang L, Qian C, Bai J, Luo W, Jin C, Yu Z. 2018. Difference in volatile composition between the pericarp tissue and inner tissue of tomato (Solanum lycopersicum) fruit. Journal of Food Processing and Preservation 42, 1–9.

Xiao H, Piovesan A, Pols S, Verboven P, Nicolaï B. 2021. Microstructural changes enhance oxygen transport in tomato (Solanum lycopersicum) fruit during maturation and ripening. New Phytologist 232, 2043–2056.

Xiao H, Verboven P, Tong S, Pedersen O, Nicolaï B. 2024. Hypoxia in tomato (Solanum lycopersicum) fruit during ripening : Biophysical elucidation by a 3D reaction–diffusion model. Plant Physiology 00, 1–13.

Ye J, Hu T, Yang C, Li H, Yang M, Ijaz R, Ye Z, Zhang Y. 2015. Transcriptome profiling of tomato fruit development reveals transcription factors associated with ascorbic acid, carotenoid and flavonoid biosynthesis. PLoS ONE 10, e0130885.

