## Supplementary for "Tissue-specific responses of the central carbon metabolism in tomato fruit to low oxygen stress"

### **Supplementary Figures**

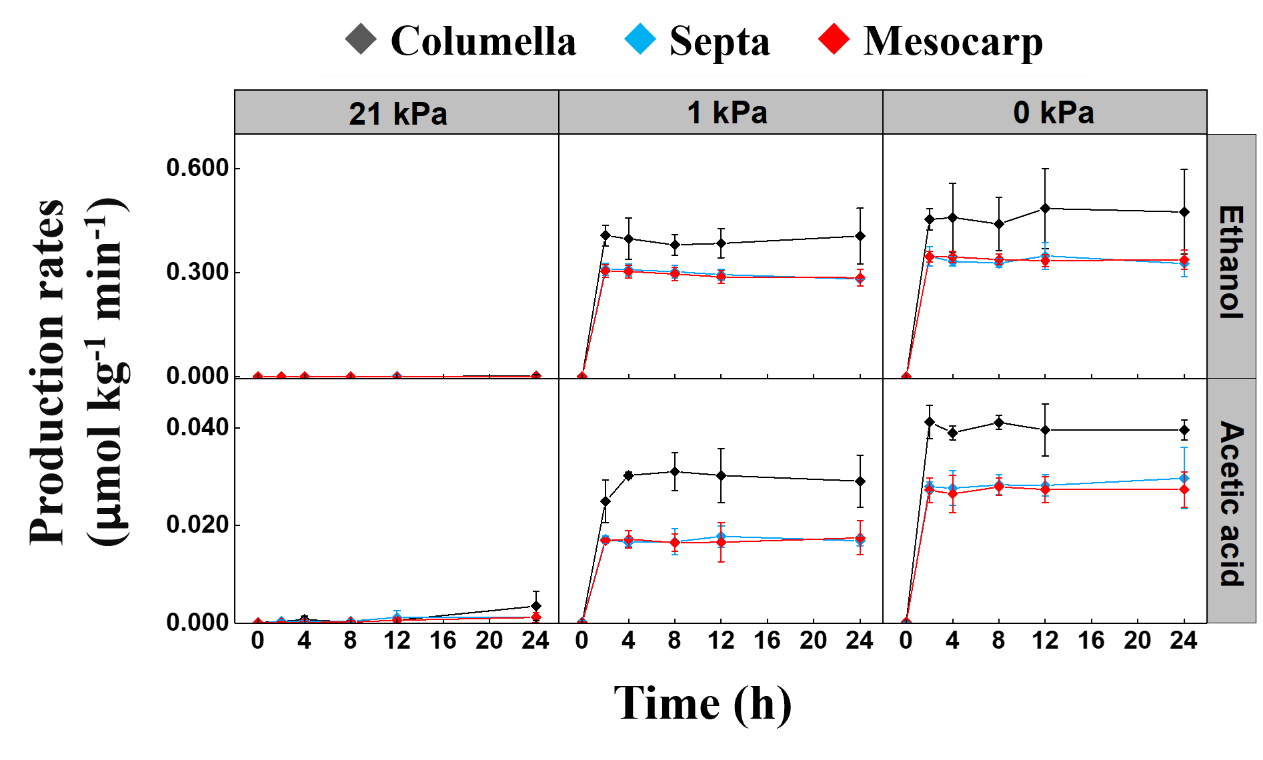

**Fig. S1.** Time series showing the production rates of ethanol and acetic acid from gas samples in columella (black), septa (blue) and mesocarp (red) tissues at 21 kPa, 1 kPa, and 0 kPa O_2_ conditions. All values represent the means of three independent replicates, and error bars indicate the standard error of the mean.

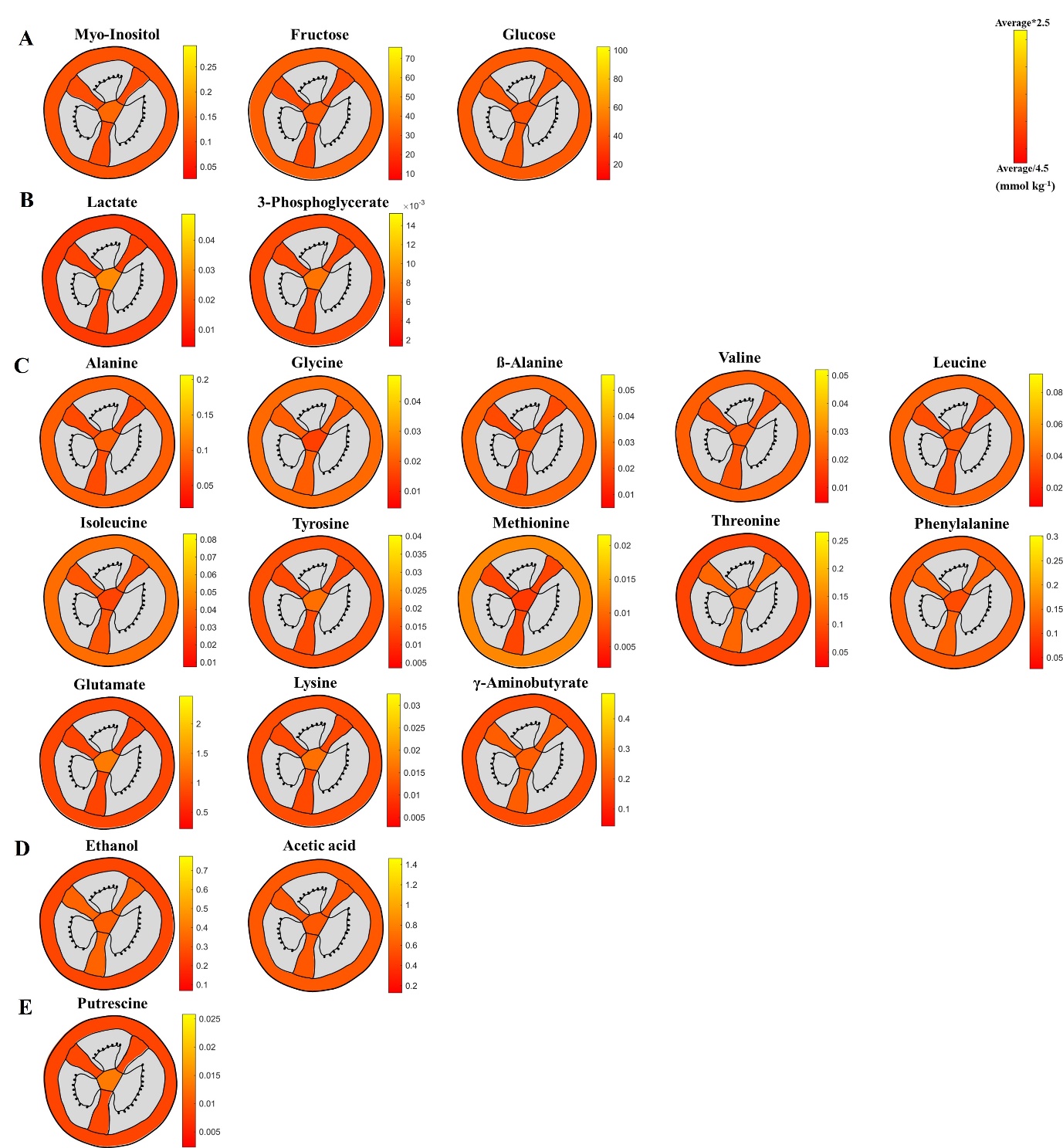

**Fig. S2.** Heat plot showing the distribution of some of the identified sugars (A), organic acids (B), amino acids (C), fermentation metabolites (D), and putrescine (E) concentration in the columella, septa, and mesocarp tissues of tomato fruit at the beginning of the incubation period (t=0 h). The scale for each plot ranges from 1/4.5 to 2.5 times the average of the metabolite concentration in the three tissues. Tissues not measured are marked in grey.

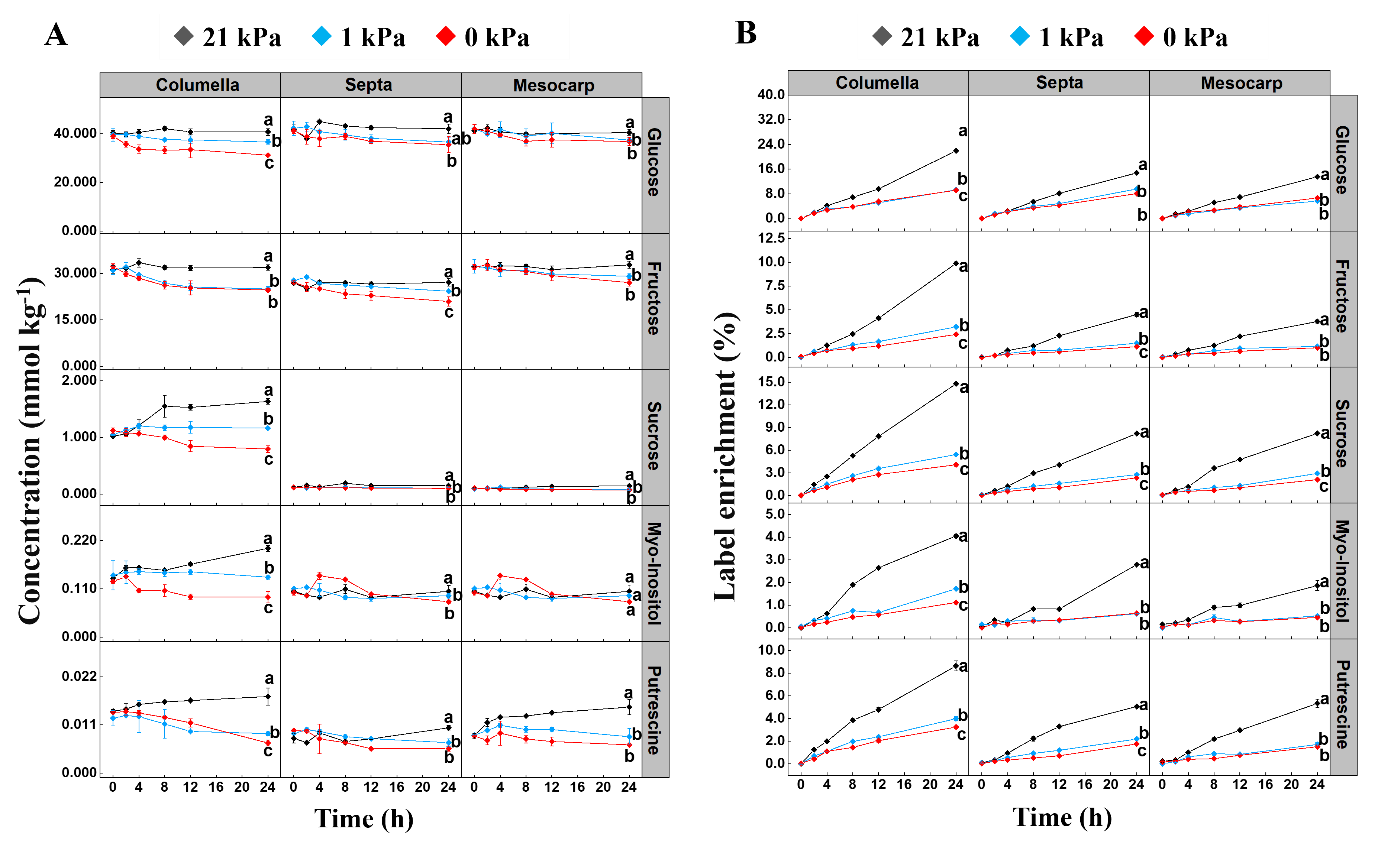

**Fig. S3.** Time series showing changes in sugar and putrescine concentrations (A) and ^13^C label enrichment (B) in three tissues after ^13^C_6_ glucose loading at 21 kPa (black), 1 kPa (blue), and 0 kPa (red) O_2_ conditions. The temperature was maintained at 18 ℃ throughout the experiment. All values represent the means of three independent replicates, and error bars indicate the standard error of the mean. Significant changes between different O_2_ concentrations in the same tissue after 24 h were based on Tukey’s honestly significant difference (HSD) test at a significant level of 0.05 and are indicated by different letters.

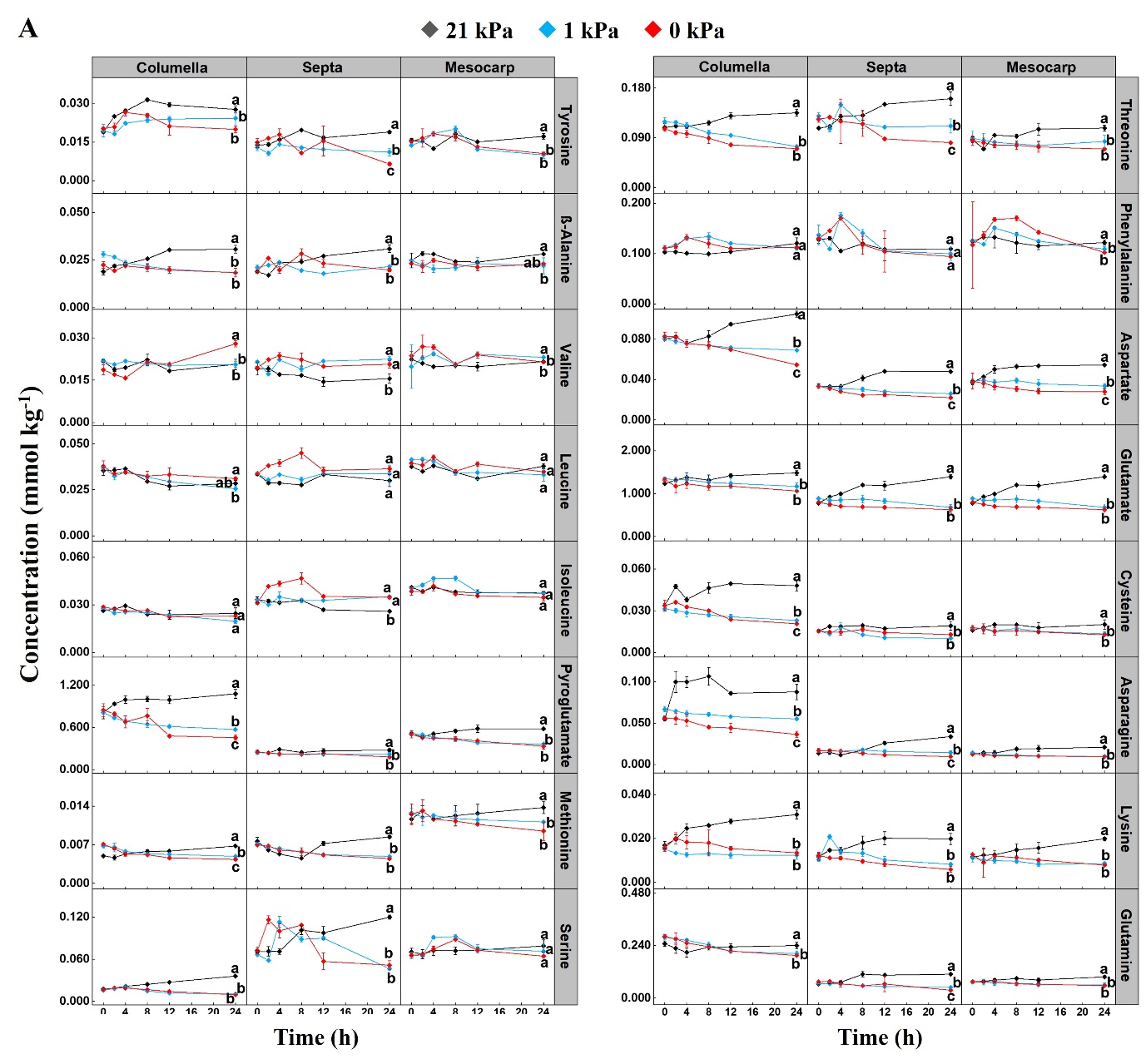

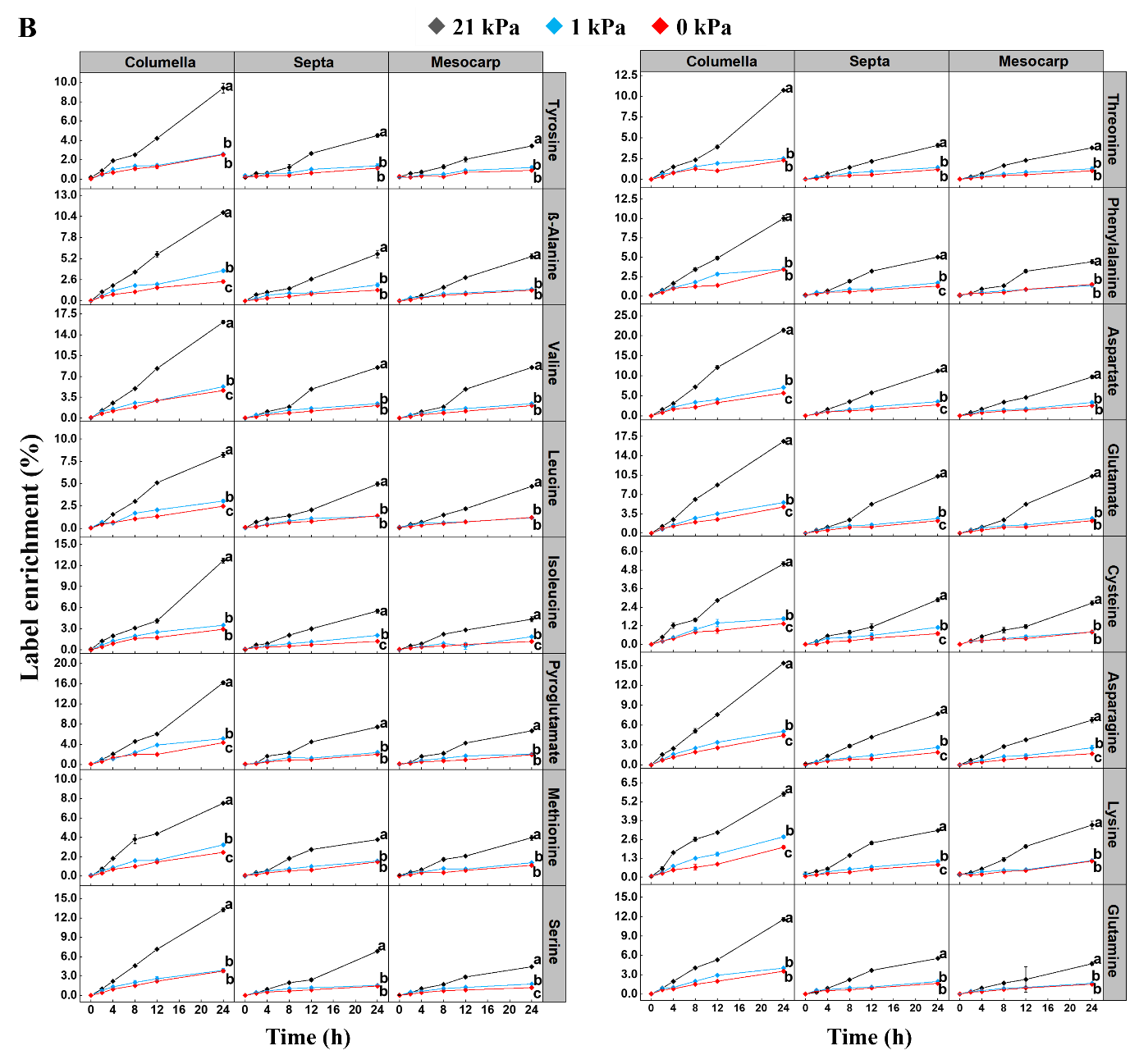

**Fig. S4.** Time series showing changes in amino acids concentrations (A) and ^13^C label enrichment (B) in three tissues after ^13^C_6_ glucose loading at 21 kPa (black), 1 kPa (blue), and 0 kPa (red) O_2_ conditions. The temperature remained constant at 18 ℃ throughout the experiment. All values represent the means of three independent replicates, and error bars indicate the standard error of the mean. Significant changes between different O_2_ concentrations in the same tissue after 24 h were based on Tukey’s honestly significant difference (HSD) test at a significant level of 0.05 and are indicated by different letters.

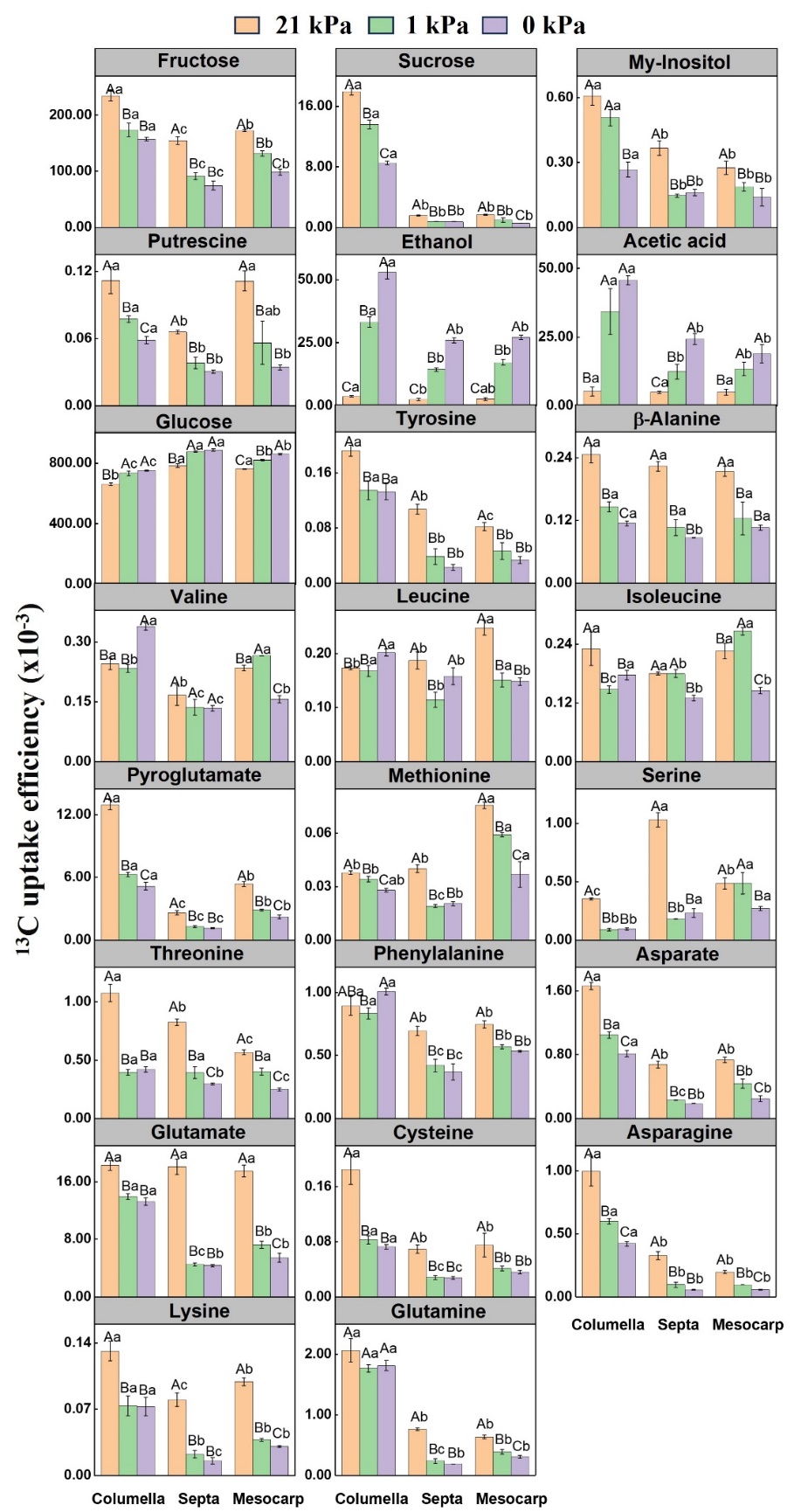

**Fig. S5.** ^13^C uptake efficiency of partial metabolites in three tissues after 24 h of incubation under three O_2_ conditions. The ^13^C uptake efficiency was calculated by multiplying the metabolite concentration by the enrichment and dividing by the total ^13^C_6_ glucose uptake equivalents. Error bars indicate the standard error of the mean. Significant changes between different O_2_ concentrations and tissues were based on Tukey’s honestly significant difference (HSD) test at a significant level of 0.05. Different capital letters show significant differences between three O_2_ conditions for the same tissue, and different lower letters indicate significant differences between three tissues in the same O_2_ condition.

### **Supplementary Tables**

**Table S1:** Osmolality of columella, septa, and mesocarp tissues. Values are the mean ± SD of nine independent measurements. Significant changes between the three tissues were based on Tukey’s honestly significant difference (HSD) test at a significant level of 0.05 and are indicated by different letters.

| **Tissue** | **Columella** | **Septa** | **Mesocarp** |
| --- | --- | --- | --- |
| Osmolality  (mOsmol kg^-1^) | 257.667±7.095a | 214±9.540b | 216±1.000b |

**Table S2:** GC-MS parameters related to the identified metabolites. Fragments and fragment-containing C-numbers were used to set the ion monitoring mode (SIM). Two different total run times indicate two different GC-MS instrumental methods.

| **Metabolites** | **Retention time (min)** | **Fragment** | **Fragment contains C number** | **Split ratio** | **Total running time (min)** | **Silylation reagent** |
| --- | --- | --- | --- | --- | --- | --- |
| Fructose | 8.47 | 189 | 2 | 1:200 | 17.3 | BSTFA |
|  |  | 364 | 4 |  |  |  |
| Glucose | 8.57 | 160 | 2 |  |  |  |
|  |  | 319 | 4 |  |  |  |
| Myo-Inositol | 9.55 | 432 | 6 | 1:20 |  |  |
| Sucrose | 12.33 | 361 | 6 |  |  |  |
|  |  | 437 | 5 |  |  |  |
| α-Ketoglutarate | 22.92 | 288 | 5 | 1:7 | 62 |  |
| Putrescine | 27.25 | 200 | 4 |  |  |  |
| 3-Phosphoglycerate | 29.4 | 459 | 3 |  |  |  |
| Fructose-6-phosphate | 41.29 | 459 | 3 |  |  |  |
| Glucose-6-phosphate | 41.52 | 160 | 2 |  |  |  |
|  |  | 471 | 4 |  |  |  |
| Lactate | 19.9 | 261 | 3 | 1:60 | 62 | MTBSTFA |
| Alanine | 21.26 | 260 | 3 |  |  |  |
| γ-Aminobutyrate | 27.28 | 274 | 4 |  |  |  |
| Serine | 33.11 | 390 | 3 |  |  |  |
| Malate | 35.89 | 419 | 4 |  |  |  |
| Aspartate | 36.83 | 418 | 4 |  |  |  |
| Glutamate | 39.35 | 432 | 5 |  |  |  |
| Citrate | 46.18 | 591 | 6 |  |  |  |
| Pyruvate | 12.68 | 174 | 3 | 1:7 | 62 |  |
| Glycine | 21.93 | 246 | 2 |  |  |  |
| ß-Alanine | 24.18 | 260 | 3 |  |  |  |
| Valine | 24.61 | 288 | 5 |  |  |  |
| Leucine | 25.75 | 302 | 6 |  |  |  |
| Isoleucine | 26.58 | 302 | 6 |  |  |  |
| Succinate | 27.39 | 289 | 4 |  |  |  |
| Fumarate | 28.15 | 287 | 4 |  |  |  |
| Pyroglutamate | 32.11 | 300 | 5 |  |  |  |
| Methionine | 32.54 | 320 | 5 |  |  |  |
| Threonine | 33.82 | 320 | 4 |  |  |  |
| Phenylalanine | 35.32 | 336 | 9 |  |  |  |
| Cysteine | 37.96 | 406 | 3 |  |  |  |
| Asparagine | 39.99 | 417 | 4 |  |  |  |
| Lysine | 41.6 | 300 | 6 |  |  |  |
| Glutamine | 42.4 | 431 | 5 |  |  |  |
| Tyrosine | 46.5 | 466 | 9 |  |  |  |

**Table S3:** Initial concentrations (mmol kg^-1^) of metabolites measured in the three tomato tissues at the beginning of the incubation period (t=0 h). Values are the mean ± SD of three independent measurements under three O_2_ conditions. Significant changes between the three tissues were based on Tukey’s honestly significant difference (HSD) test at a significant level of 0.05 and are indicated by different letters. The fold changes in the concentration of each compound in the three tissues were determined by comparing it with the lowest concentration value for each compound among the three tissues.

|  |  | **Concentration (mmol kg^-1^)** | | | **Fold change** | | |
| --- | --- | --- | --- | --- | --- | --- | --- |
|  |  | **Columella** | **Septa** | **Mesocarp** | **Columella** | **Septa** | **Mesocarp** |
| **Sugars** | Fructose-6-phosphate | 0.0905±0.0018a | 0.0354±0.0031c | 0.0438±0.0041b | 2.5565 | 1.0000 | 1.2373 |
|  | Glucose-6-phosphate | 0.1072±0.0107a | 0.0393±0.0078b | 0.0392±0.0043b | 2.7347 | 1.0026 | 1.0000 |
|  | Fructose | 31.5636±1.2186a | 27.2968±0.5098b | 32.2162±1.2102a | 1.1563 | 1.0000 | 1.1802 |
|  | Glucose | 39.6603±1.6088b | 41.6749±2.0241a | 41.7043±1.3638a | 1.0000 | 1.0508 | 1.0515 |
|  | Myo-Inositol | 0.1341±0.0182a | 0.1056±0.0056b | 0.1119±0.0118b | 1.2699 | 1.0000 | 1.0597 |
|  | Sucrose | 1.0641±0.0495a | 0.119±0.005b | 0.1002±0.0127b | 10.6198 | 1.1876 | 1.0000 |
| **Organic acids** | Pyruvate | 0.0441±0.0047a | 0.0235±0.0036b | 0.0177±0.0009c | 2.4915 | 1.3277 | 1.0000 |
|  | Succinate | 0.0258±0.0012a | 0.0095±0.0007b | 0.0101±0.0008b | 2.7158 | 1.0000 | 1.0632 |
|  | Fumarate | 0.2663±0.0189a | 0.0896±0.0102b | 0.1176±0.0363b | 2.9721 | 1.0000 | 1.3125 |
|  | Malate | 1.4516±0.0344b | 0.6853±0.0222c | 1.6386±0.1366a | 2.1182 | 1.0000 | 2.3911 |
|  | Citrate | 0.7592±0.0361b | 2.0993±0.1168a | 2.1894±0.173a | 1.0000 | 2.7651 | 2.8838 |
|  | α-Ketoglutarate | 0.2698±0.0138a | 0.0966±0.009b | 0.0995±0.0128b | 2.7930 | 1.0000 | 1.0300 |
|  | Lactate | 0.0281±0.0024a | 0.0159±0.0017b | 0.0145±0.0022b | 1.9379 | 1.0966 | 1.0000 |
|  | 3-Phosphoglycerate | 0.0077±0.0004a | 0.0054±0.0005b | 0.0053±0.0002b | 1.4528 | 1.0189 | 1.0000 |
| **Amino acids** | Alanine | 0.088±0.0047a | 0.073±0.0056b | 0.0868±0.0081a | 1.2055 | 1.0000 | 1.1890 |
|  | γ-Aminobutyrate | 0.2044±0.0127a | 0.2057±0.013a | 0.1709±0.0077b | 1.1960 | 1.2036 | 1.0000 |
|  | Glycine | 0.0153±0.0013c | 0.0206±0.0012b | 0.0226±0.0013a | 1.0000 | 1.3464 | 1.4771 |
|  | ß-Alanine | 0.0231±0.0042a | 0.0197±0.0013b | 0.0242±0.0021a | 1.1726 | 1.0000 | 1.2284 |
|  | Valine | 0.0207±0.0018a | 0.02±0.0017a | 0.0219±0.0043a | 1.0350 | 1.0000 | 1.0950 |
|  | Leucine | 0.0367±0.0023b | 0.0337±0.0006c | 0.0395±0.0017a | 1.0890 | 1.0000 | 1.1721 |
|  | Isoleucine | 0.0275±0.001c | 0.0325±0.0014b | 0.04±0.0017a | 1.0000 | 1.1818 | 1.4545 |
|  | Pyroglutamate | 0.823±0.0667a | 0.2465±0.0189c | 0.5112±0.0332b | 3.3387 | 1.0000 | 2.0738 |
|  | Methionine | 0.0063±0.001b | 0.0073±0.0005b | 0.0123±0.0012a | 1.0000 | 1.1587 | 1.9524 |
|  | Serine | 0.0167±0.0011b | 0.0699±0.0036a | 0.0671±0.0045a | 1.0000 | 4.1856 | 4.0180 |
|  | Threonine | 0.1116±0.0081a | 0.1203±0.0106a | 0.0871±0.0082b | 1.2809 | 1.3815 | 1.0000 |
|  | Phenylalanine | 0.1081±0.005a | 0.1315±0.0113a | 0.1225±0.043a | 1.0000 | 1.2165 | 1.1332 |
|  | Aspartate | 0.0813±0.0028a | 0.0335±0.0016c | 0.0381±0.0043b | 2.4267 | 1.0000 | 1.1367 |
|  | Cysteine | 0.0329±0.0025a | 0.0156±0.0008b | 0.0173±0.0012b | 2.1090 | 1.0000 | 1.1090 |
|  | Glutamate | 1.3007±0.0621a | 0.8218±0.0539b | 0.8442±0.0505b | 1.5827 | 1.0000 | 1.0273 |
|  | Asparagine | 0.0594±0.0061a | 0.0158±0.0023b | 0.0133±0.0008b | 4.4662 | 1.1880 | 1.0000 |
|  | Lysine | 0.0159±0.0015a | 0.0114±0.0015b | 0.0118±0.0013b | 1.3947 | 1.0000 | 1.0351 |
|  | Glutamine | 0.2696±0.0178a | 0.0693±0.0055b | 0.0750±0.0060b | 3.8885 | 1.0000 | 1.0812 |
|  | Tyrosine | 0.0195±0.0014a | 0.0139±0.0014b | 0.0149±0.001b | 1.4029 | 1.0000 | 1.0719 |
| **Fermentation metabolites** | Ethanol | 0.3245±0.0305ab | 0.3417±0.078a | 0.2628±0.054b | 1.2348 | 1.3002 | 1.0000 |
|  | Acetic acid | 0.5869±0.168a | 0.5788±0.1415a | 0.5884±0.1122a | 1.0140 | 1.0000 | 1.0166 |
| **Polyamine** | Putrescine | 0.0135±0.0012a | 0.0089±0.001b | 0.0086±0.0003b | 1.5698 | 1.0349 | 1.0000 |

**Table S4:** Estimated Michaelis-Menten parameters for O_2_ consumption and CO_2_ production rates. The standard errors are given in brackets.

| **Tissue specific parameters:** | **Tissue** | | | |
| --- | --- | --- | --- | --- |
|  | **Columella** | | **Septa** | **Mesocarp** |
| *V*_max,O2_ | | 1193.50 (41.92) | 612.59 (24.25) | 519.93 (21.90) |
| *V*_max,CO2,_ | | 524.09 (20.90) | 266.05 (17.50) | 210.04 (17.38) |
| **Parameters estimated in common:** | | | | |
| *K*_m,O2_ | | 7.2007 (0.6354) | | |
| *RQ* | | 0.9104 (0.0210) | | |
| *K*_m,f,O2_ | | 0.6293 (0.1025) | | |

*V*_max,O2_ (nmol kg^-1^ s^-1^) is the maximal O_2_ consumption rate; *RQ* is the respiratory quotient; *V*_max,CO2_ (nmol kg^-1^ s^-1^) is the maximal fermentative CO_2_ production rate; *K*_m,O2_ is the Michaelis constant for O_2_ consumption (kPa); *K*_m,f,O2_ is the Michaelis constant for the competitive inhibition of fermentative CO_2_ production by O_2_ (kPa).

**Table S5:** Independent t-test p-values were calculated to compare metabolite concentrations measured at the beginning (t=0 h) and end (t=24 h) of the incubation period for the three tissues under 21, 1, and 0 kPa O_2_. P-values greater than 0.05 indicate that the difference is not significant, and are marked in grey. P-values less than 0.05 are marked in red to indicate that the concentration at 24 h was significantly higher than that at 0 h of the incubation period. Conversely, p-values less than 0.05 are marked in green to indicate that the concentration at 0 h was significantly higher than that at 24 h of the incubation period.

|  |  | **21 kPa** | | | **1 kPa** | | | **0 kPa** | | |
| --- | --- | --- | --- | --- | --- | --- | --- | --- | --- | --- |
|  |  | **Columella** | **Septa** | **Mesocarp** | **Columella** | **Septa** | **Mesocarp** | **Columella** | **Septa** | **Mesocarp** |
| **Sugars** | Fructose-6-phosphate | 0 | 0.046 | 0.162 | 0.003 | 0 | 0.001 | 0 | 0.003 | 0.004 |
|  | Glucose-6-phosphate | 0.022 | 0.592 | 0.656 | 0.006 | 0.014 | 0.32 | 0 | 0 | 0.03 |
|  | Fructose | 0.559 | 0.752 | 0.459 | 0.003 | 0 | 0.078 | 0 | 0.002 | 0.003 |
|  | Glucose | 0.706 | 0.827 | 0.397 | 0.134 | 0.046 | 0.014 | 0 | 0.046 | 0.023 |
|  | Myo-Inositol | 0.002 | 1 | 0.779 | 0.844 | 0.011 | 0.028 | 0.01 | 0.007 | 0.236 |
|  | Sucrose | 0.002 | 0.004 | 0.001 | 0 | 0.016 | 0.416 | 0.002 | 0.012 | 0.129 |
| **Organic acids** | Pyruvate | 0.269 | 0.1 | 0.483 | 0.026 | 0.009 | 0 | 0.006 | 0.003 | 0.021 |
|  | Succinate | 0 | 0 | 0.018 | 0.001 | 0.188 | 0 | 0 | 0 | 0 |
|  | Fumarate | 0.003 | 0.016 | 0.219 | 0.001 | 0.003 | 0 | 0 | 0 | 0.142 |
|  | Malate | 0.001 | 0 | 0.001 | 0 | 0.004 | 0 | 0 | 0.025 | 0.014 |
|  | Citrate | 0.001 | 0.004 | 0.017 | 0.014 | 0 | 0.071 | 0.006 | 0.001 | 0.008 |
|  | α-Ketoglutarate | 0 | 0.075 | 0.043 | 0.006 | 0.029 | 0.225 | 0 | 0.005 | 0.003 |
|  | Lactate | 0.05 | 0.794 | 0.046 | 0 | 0.001 | 0 | 0 | 0.006 | 0 |
|  | 3-Phosphoglycerate | 0.341 | 0.947 | 0.049 | 0.575 | 0.001 | 0.005 | 0.003 | 0.001 | 0 |
| **Amino acids** | Alanine | 0 | 0 | 0.001 | 0 | 0 | 0 | 0 | 0 | 0 |
|  | γ-Aminobutyrate | 0.002 | 0.823 | 0.781 | 0 | 0.02 | 0.009 | 0.003 | 0.015 | 0.023 |
|  | Glycine | 0.09 | 0.049 | 0.331 | 0.012 | 0.863 | 0 | 0.002 | 0.006 | 0.001 |
|  | ß-Alanine | 0.002 | 0.006 | 0.262 | 0 | 0.518 | 0.378 | 0.068 | 0.374 | 0.431 |
|  | Valine | 0.399 | 0.1 | 0.015 | 0.283 | 0.193 | 0.54 | 0.002 | 0.197 | 0.111 |
|  | Leucine | 0.016 | 0.145 | 0.732 | 0 | 0.994 | 0.054 | 0.017 | 0.04 | 0 |
|  | Isoleucine | 0.42 | 0.003 | 0.002 | 0 | 0.013 | 0.026 | 0 | 0.009 | 0.109 |
|  | Pyroglutamate | 0.013 | 0.208 | 0.012 | 0 | 0.323 | 0.001 | 0.002 | 0 | 0.011 |
|  | Methionine | 0 | 0.139 | 0.053 | 0.001 | 0 | 0.033 | 0 | 0.001 | 0.103 |
|  | Serine | 0 | 0 | 0.222 | 0 | 0 | 0.391 | 0.001 | 0.01 | 0.522 |
|  | Threonine | 0.008 | 0.002 | 0.008 | 0.001 | 0.089 | 0.567 | 0 | 0 | 0.09 |
|  | Phenylalanine | 0.057 | 0.011 | 0.441 | 0.93 | 0.052 | 0.026 | 0.677 | 0 | 0.776 |
|  | Aspartate | 0 | 0.001 | 0 | 0.001 | 0 | 0.061 | 0.006 | 0.001 | 0.091 |
|  | Cysteine | 0.004 | 0.173 | 0.172 | 0 | 0.001 | 0.009 | 0.028 | 0.048 | 0.006 |
|  | Glutamate | 0.002 | 0 | 0.001 | 0.026 | 0.008 | 0.005 | 0.002 | 0.042 | 0.107 |
|  | Asparagine | 0.004 | 0 | 0.011 | 0.005 | 0.213 | 0.031 | 0.001 | 0.011 | 0.002 |
|  | Lysine | 0.001 | 0.01 | 0 | 0.089 | 0.035 | 0.091 | 0.129 | 0.006 | 0 |
|  | Glutamine | 0.502 | 0.003 | 0.013 | 0.001 | 0.014 | 0.113 | 0 | 0 | 0.014 |
|  | Tyrosine | 0 | 0.001 | 0.113 | 0.089 | 0.184 | 0.01 | 0.715 | 0.001 | 0 |
| **Fermentation metabolites** | Ethanol | 0.14 | 0.392 | 0.098 | 0.002 | 0 | 0 | 0.001 | 0 | 0.001 |
|  | Acetic acid | 0.857 | 0.077 | 0.885 | 0.004 | 0.003 | 0.021 | 0 | 0 | 0.002 |
| **Polyamine** | Putrescine | 0.044 | 0.024 | 0.004 | 0.03 | 0.018 | 0.762 | 0 | 0 | 0 |

**Table S6:** Enrichment fractions (%) of all labeled metabolites in the three tomato tissues at 24 h of ^13^C_6_ glucose incubation at 21 kPa. Values are the mean ± SD of three independent measurements. Significant changes were based on Tukey’s honestly significant difference (HSD) test at a significant level of 0.05 and indicated by different letters.

|  |  | **Columella** | **Septa** | **Mesocarp** |
| --- | --- | --- | --- | --- |
| **Sugars** | Fructose-6-phosphate | 15.3642±0.4588a | 7.3029±0.2608b | 7.687±2.047b |
|  | Glucose-6-phosphate | 21.1665±0.3507a | 9.4054±0.8242b | 9.0973±0.4001b |
|  | Fructose | 9.8917±0.1618a | 4.522±0.2231b | 3.7893±0.1762c |
|  | Glucose | 21.9467±0.1382a | 14.7971±0.2365b | 13.567±0.0947c |
|  | Myo-Inositol | 4.0508±0.0824a | 2.7899±0.0376b | 1.8594±0.2141c |
|  | Sucrose | 14.7947±0.1707a | 8.2009±0.0711b | 8.2303±0.1164b |
| **Organic acids** | Pyruvate | 12.9101±0.2173a | 6.1826±0.1232b | 5.3901±0.0824c |
|  | Succinate | 13.8105±0.1225a | 6.238±0.3609b | 6.6256±0.1981b |
|  | Fumarate | 12.5451±0.0672a | 6.7671±0.1055b | 6.3473±0.0897c |
|  | Malate | 9.589±0.4788a | 5.0528±0.1918b | 3.6595±0.0622c |
|  | Citrate | 12.4011±0.3369a | 7.1704±0.2538b | 5.6088±0.2802c |
|  | α-Ketoglutarate | 15.8433±0.2882a | 7.4009±0.3256b | 7.288±0.1683b |
|  | Lactate | 2.3516±0.0476a | 1.5036±0.2159b | 1.1018±0.1748b |
|  | 3-Phosphoglycerate | 13.4356±0.249a | 5.9235±0.1473c | 6.7579±0.254b |
| **Amino acids** | Alanine | 13.1713±0.1022a | 6.497±0.0834b | 5.4951±0.0609c |
|  | γ-Aminobutyrate | 10.8167±0.087a | 5.6873±0.1279b | 5.0972±0.2289c |
|  | Glycine | 8.696±0.184a | 4.2613±0.1839b | 3.7228±0.0437c |
|  | ß-Alanine | 10.8696±0.0716a | 5.7651±0.4553b | 5.4691±0.2948b |
|  | Valine | 16.109±0.2822a | 8.5044±0.0644b | 7.8263±0.1287c |
|  | Leucine | 8.2178±0.2826a | 4.9599±0.2578b | 4.696±0.0346b |
|  | Isoleucine | 12.6809±0.3593a | 5.5239±0.265b | 4.3562±0.3439c |
|  | Pyroglutamate | 16.2081±0.3344a | 7.4382±0.2242b | 6.6682±0.1735c |
|  | Methionine | 7.519±0.0223a | 3.7681±0.0479b | 3.9618±0.262b |
|  | Serine | 13.2554±0.2645a | 6.8359±0.1915b | 4.4354±0.1636c |
|  | Threonine | 10.7324±0.0143a | 4.0845±0.2445b | 3.7904±0.1086b |
|  | Phenylalanine | 10.023±0.3674a | 5.0288±0.0231b | 4.412±0.2614b |
|  | Aspartate | 21.3921±0.5142a | 11.2151±0.2567b | 9.7036±0.4685c |
|  | Cysteine | 5.1945±0.1436a | 2.8784±0.123b | 2.6636±0.148b |
|  | Glutamate | 16.6129±0.2332a | 10.283±0.1957b | 7.7162±0.2382c |
|  | Asparagine | 15.3284±0.1773a | 7.6907±0.2003b | 6.7501±0.4094c |
|  | Lysine | 5.7334±0.1419a | 3.2097±0.07b | 3.5908±0.2668b |
|  | Glutamine | 11.5731±0.2737a | 5.5278±0.101b | 4.7063±0.2442c |
|  | Tyrosine | 9.3977±0.4888a | 4.5063±0.1731b | 3.4316±0.1212c |
| **Polyamine** | Putrescine | 8.6416±0.4524a | 5.0587±0.0588b | 5.3307±0.3777b |

**Table S7:** Enrichment fractions (%) of all labeled metabolites in the three tomato tissues at 24 h of ^13^C_6_ glucose incubation at 1 kPa. Values are the mean ± SD of three independent measurements. Significant changes were based on Tukey’s honestly significant difference (HSD) test at a significant level of 0.05 and indicated by different letters.

|  |  | **Columella** | **Septa** | **Mesocarp** |
| --- | --- | --- | --- | --- |
| **Sugars** | Fructose-6-phosphate | 7.8898±0.2016a | 3.9442±0.2439b | 3.6193±0.2664b |
|  | Glucose-6-phosphate | 9.5939±0.0688a | 5.21±0.0125b | 4.0758±0.2849c |
|  | Fructose | 3.2293±0.132a | 1.5101±0.0859b | 1.1741±0.0928c |
|  | Glucose | 9.3282±0.0636a | 9.6105±0.212a | 5.6907±0.0266b |
|  | Myo-Inositol | 1.7186±0.0491a | 0.6208±0.0137b | 0.5259±0.044c |
|  | Sucrose | 5.4232±0.075a | 2.7734±0.1596b | 2.9322±0.1571b |
| **Organic acids** | Pyruvate | 5.6847±0.1956a | 2.7571±0.1916b | 2.462±0.0735b |
|  | Succinate | 6.2234±0.3216a | 2.7302±0.1995b | 2.8262±0.0751b |
|  | Fumarate | 4.1561±0.102a | 2.4321±0.1367b | 2.0286±0.0837c |
|  | Malate | 2.3948±0.096a | 1.5838±0.0708b | 1.0197±0.077c |
|  | Citrate | 4.6991±0.0522a | 2.2413±0.0801b | 1.8718±0.2945b |
|  | α-Ketoglutarate | 5.1802±0.1408a | 2.5132±0.2036b | 2.6065±0.2692b |
|  | Lactate | 6.8074±0.6019a | 3.7598±0.1554b | 3.0199±0.1335c |
|  | 3-Phosphoglycerate | 5.9333±0.7295a | 2.9131±0.1061b | 2.857±0.0087c |
| **Amino acids** | Alanine | 5.7542±0.0842a | 2.8432±0.0588b | 2.271±0.0994c |
|  | γ-Aminobutyrate | 4.8435±0.0716a | 2.1329±0.1063b | 2.4277±0.2923b |
|  | Glycine | 3.7778±0.1145a | 1.7348±0.1973b | 1.6576±0.0818b |
|  | ß-Alanine | 3.7169±0.1827a | 1.9625±0.2354b | 1.4417±0.1235c |
|  | Valine | 5.2912±0.0884a | 2.4271±0.1961c | 2.9828±0.0087b |
|  | Leucine | 3.0536±0.1627a | 1.3599±0.0953b | 1.1798±0.0509b |
|  | Isoleucine | 3.4938±0.0987a | 2.0557±0.0765b | 1.8623±0.0406c |
|  | Pyroglutamate | 5.1132±0.0696a | 2.3747±0.0881b | 2.04±0.047c |
|  | Methionine | 3.22±0.1213a | 1.5515±0.0516b | 1.3724±0.0246b |
|  | Serine | 3.8813±0.3378a | 1.5451±0.1111b | 1.7612±0.1245b |
|  | Threonine | 2.4863±0.1435a | 1.4176±0.1097b | 1.2692±0.2292b |
|  | Phenylalanine | 3.4938±0.2035a | 1.6937±0.0632b | 1.3555±0.1044b |
|  | Aspartate | 7.0503±0.1695a | 3.5386±0.1868b | 3.3374±0.3369b |
|  | Cysteine | 1.6594±0.1037a | 1.0969±0.0953b | 0.7979±0.0632c |
|  | Glutamate | 5.538±0.1031a | 2.6551±0.1007b | 2.4765±0.0923b |
|  | Asparagine | 5.0514±0.0642a | 2.6384±0.2191b | 2.5705±0.3946b |
|  | Lysine | 2.779±0.0946a | 1.081±0.1098b | 1.1412±0.106b |
|  | Glutamine | 4.0272±0.0887a | 1.9485±0.047b | 1.674±0.11c |
|  | Tyrosine | 2.5897±0.0097a | 1.3638±0.2292b | 1.1978±0.1718b |
| **Polyamine** | Putrescine | 3.9892±0.17a | 2.1955±0.0495b | 1.7149±0.1808c |

**Table S8:** Enrichment fractions (%) of all labeled metabolites in the three tomato tissues at 24 h of ^13^C_6_ glucose incubation at 0 kPa. Values are the mean ± SD of three independent measurements. Significant changes were based on Tukey’s honestly significant difference (HSD) test at a significant level of 0.05 and indicated by different letters.

|  |  | **Columella** | **Septa** | **Mesocarp** |
| --- | --- | --- | --- | --- |
| **Sugars** | Fructose-6-phosphate | 5.7015±0.1425a | 2.7535±0.1751b | 2.3485±0.1072c |
|  | Glucose-6-phosphate | 7.6653±0.1355a | 3.8138±0.0398b | 3.5611±0.0627b |
|  | Fructose | 2.4302±0.0924a | 1.1469±0.0287b | 1.0345±0.0368c |
|  | Glucose | 9.1959±0.0409a | 8.1155±0.3675b | 6.6891±0.1423c |
|  | Myo-Inositol | 1.1161±0.0716a | 0.6432±0.0348b | 0.4571±0.0513c |
|  | Sucrose | 4.0701±0.215a | 2.3359±0.0095b | 2.1074±0.0185b |
| **Organic acids** | Pyruvate | 4.6311±0.112a | 2.4533±0.0647b | 2.1005±0.0302c |
|  | Succinate | 4.7377±0.1355a | 2.4637±0.1207b | 2.3026±0.0126b |
|  | Fumarate | 3.2915±0.1474a | 1.7086±0.0396b | 1.6142±0.0342b |
|  | Malate | 2.281±0.1133a | 0.7666±0.0539b | 0.9114±0.0413b |
|  | Citrate | 3.6506±0.146a | 1.7445±0.0945b | 1.5132±0.0361b |
|  | α-Ketoglutarate | 4.4542±0.1966a | 2.1392±0.0668b | 2.0964±0.0217b |
|  | Lactate | 8.933±0.3304a | 4.6831±0.4858b | 3.6235±0.4308c |
|  | 3-Phosphoglycerate | 4.6541±0.126a | 2.4534±0.0924b | 2.2304±0.3105b |
| **Amino acids** | Alanine | 4.8296±0.071a | 2.3796±0.0401b | 2.1875±0.0945c |
|  | γ-Aminobutyrate | 3.9175±0.1938a | 2.0825±0.1855b | 1.8699±0.0215b |
|  | Glycine | 3.3077±0.1298a | 1.5341±0.0742b | 1.4391±0.0061b |
|  | ß-Alanine | 2.3795±0.1799a | 1.3222±0.08b | 1.3161±0.0309b |
|  | Valine | 4.6401±0.1969a | 2.0976±0.0487b | 2.0795±0.0769b |
|  | Leucine | 2.4741±0.0872a | 1.4002±0.0585b | 1.2125±0.0472c |
|  | Isoleucine | 2.9414±0.1144a | 1.2125±0.0571b | 1.1986±0.04b |
|  | Pyroglutamate | 4.3241±0.074a | 2.0314±0.0073b | 1.9069±0.0156c |
|  | Methionine | 2.4466±0.0981a | 1.4632±0.0262b | 1.1088±0.0731c |
|  | Serine | 3.7677±0.212a | 1.4498±0.0476b | 1.2019±0.0637b |
|  | Threonine | 2.2776±0.034a | 1.1779±0.0139b | 1.0226±0.0313c |
|  | Phenylalanine | 3.419±0.0902a | 1.2683±0.2217b | 1.4873±0.0329b |
|  | Aspartate | 5.6536±0.1562a | 2.7213±0.0618b | 2.5178±0.0631b |
|  | Cysteine | 1.3397±0.0424a | 0.6933±0.1015b | 0.7996±0.0771b |
|  | Glutamate | 4.7344±0.1664a | 2.2561±0.1573b | 2.1642±0.0442b |
|  | Asparagine | 4.4276±0.2986a | 1.8877±0.0947b | 1.696±0.0999b |
|  | Lysine | 2.0545±0.1116a | 0.8587±0.0268c | 1.1074±0.0756b |
|  | Glutamine | 3.5396±0.1763a | 1.6642±0.0292b | 1.5349±0.0443b |
|  | Tyrosine | 2.5366±0.1501a | 1.1327±0.12b | 0.8992±0.1278b |
| **Polyamine** | Putrescine | 3.2487±0.1127a | 1.7624±0.0561b | 1.5066±0.0377c |

#### Supplementary Method S1

**Determination of the preincubation time**

A preincubation stage was implemented to allow tissues to recover from handling damage and adapt to experimental conditions, stabilizing their metabolic state under 21 kPa O_2_ at 18 ℃. To determine the optimal preincubation duration, the O₂ consumption rate of the three tissues in 20 mmol L^-1^ glucose buffer was measured at various time points using the OX1LP dissolved O_2_ package (Qubit Systems, Kingston, Ontario, Canada), which employs polarographic dissolved O_2_ electrodes. Before measurement, 3 mL of buffer solution was added to each chamber, with calibration performed at two known levels: zero O₂ concentration and air-saturated levels. After calibration, tissue samples were added to the air-saturated buffer, and O₂ consumption was recorded for approximately 30 min per sample.

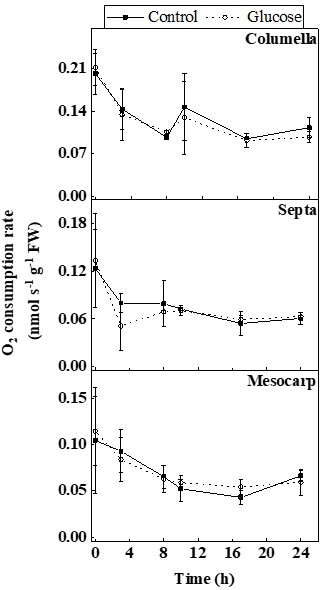

**Fig. SM1.** Changes in O₂ consumption rate of three tomato fruit tissues after incubation in empty buffer without glucose (control) and 20 mmol L^-1^ glucose buffer at 18 °C and 21 kPa O_2_^.^ Error bars represent the standard error of the mean.

The results showed that the O₂ consumption rate decreased from 0 to 17 h of incubation and stabilized between 17 and 24 h. Based on these findings, a 24-h preincubation period was selected. The similar O₂ consumption rates between the control and glucose buffer over 24 h confirmed that the 20 mmol L^-1^ glucose buffer does not alter tissue respiration.
